# Mapping the GDF15 Arm of the Integrated Stress Response in Human Cells and Tissues

**DOI:** 10.1101/2025.01.31.635929

**Authors:** Janell L.M. Smith, Kamaryn Tanner, Jack Devine, Anna S. Monzel, Taivan Batjargal, Maxwell Z. Wilson, Alan A. Cohen, Martin Picard

## Abstract

Mitochondrial stress activates the integrated stress response (ISR) and triggers cell–cell communication through the secretion of the metabokine growth differentiation factor 15 (GDF15). However, the gene network underlying the ISR remains poorly defined, particularly across metabolically diverse cellular states and tissues. Using RNAseq data from fibroblasts subjected to eleven metabolic perturbations, including genetic and pharmacological mitochondrial OxPhos defects, we showed that the ISR has multiple arms and developed an *ISR^GDF15^* index quantifying the GDF15 arm of ISR activation in human cells. The *ISR^GDF15^* index was validated using optogenetic activation of the ISR protein kinase R (PKR) in a stable cell line, demonstrating its rapid kinetics preceding to *GDF15* gene expression. We then deployed the *ISR^GDF15^* index across 44 postmortem human tissues, reporting that the *ISR^GDF15^* was upregulated in the heart of individuals who died of an acute cause in the emergency room, whereas it was preferentially upregulated in the brain of individuals who died as inpatients after protracted hospital stays. *ISR^GDF15^* was also moderately, positively correlated with age across all tissues. These data highlight multiple distinct ISR pathways and clarify which genes are related to the GDF15 arm of the ISR, yielding an *ISR^GDF15^* index that can be used to investigate tissue-specific and age-related ISR activation in both in vitro cultures and human tissues. Ultimately, we expand our knowledge of the GDF15 arm of the ISR in humans, show its inducibility by the ISR kinase PKR, and demonstrate its applicability to detect ISR activation in specific human tissues.

## Main Text

Cells sense reductive stress via the integrated stress response (ISR)^1^. Among many targets, the ISR activates the gene *GDF15*, which encodes a secreted cytokine/metabokine known to be elevated with aging^2^ and in several chronic diseases^3,4^. This elevation is particularly strong in primary mitochondrial diseases caused by defects in the oxidative phosphorylation (OxPhos) system^5,6^. However, the ISR is pleiotropic and triggers multiple cellular processes^7^. As a result, there is no consensus on the gene network that constitutes the ISR in human cells, or whether the same set of genes operate in metabolically diverse organs or tissues. Here, we seek to understand the dimensionality of the ISR, with a particular focus on the pathway or “arm” leading to GDF15 signaling, given its status as an ISR biomarker^8^ and its role in mediating the ISR^9^.

To identify ISR-related genes linked to inter-cellular GDF15 signaling, we queried multiple ISR gene lists (119 genes in total)^10–12^ (**Figure 1A**), and examined their co-expression in a cellular-lifespan-system dataset generated in fibroblasts^13^. Sensitivity analyses were performed on each gene list separately, as well as a random selection of 119 selected genes out of all mapped genes in the RNAseq dataset. This process demonstrated the superiority of including all 119 genes from the combined lists, plus *GDF15*, in the fibroblast dataset, over using the individual ISR gene lists, equally loaded genes of the 119 combined genes, or 119 randomly chosen genes (**Extended Data Figure 1**).

**Figure 1.**
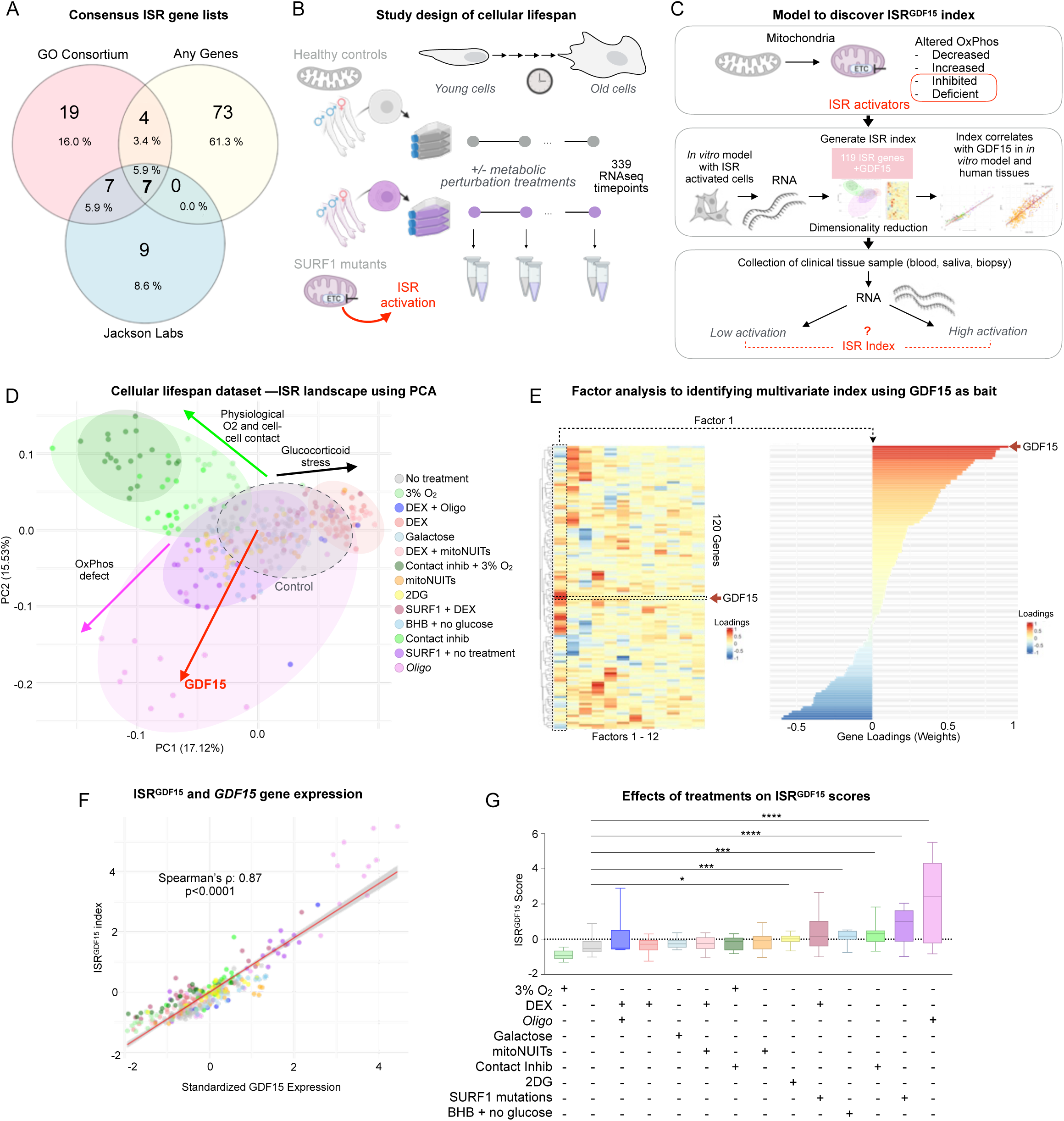
Development of an ISR index using primary human fibroblasts. (**A**) ISR related genes from three reference ISR pathway lists revealing minimal overlap. We use all 119 genes plus *GDF15* (n=120) to compile an unbiased ISR gene list. (**B**) Bulk RNA seq data in primary human fibroblasts from three healthy donors and three individuals with mitochondrial disease (SURF1 gene mutations) were cultured across their replicative lifespans (up to 260 days) and treated with pharmacological and metabolic perturbations, including the OxPhos inhibitor oligomycin (1nM). (**C**) Overview of the analytical strategy to generate an ISR index using *GDF15* as a bait. (**D**) Principal component analysis (PCA) of data from the lifespan study depicted in panel B including 339 samples 120 genes. The red arrow denotes the loading vector of *GDF15*; all other genes shown in **Extended Data Figure 2**. (**E**) Factor analysis revealed one factor where *GDF15* had the strongest positive loading (Factor 1, *left*), yielding weights that reflect the co-regulation of all other genes with *GDF15* (*right*). (**F**) Correlation of Factor 1 with standardized *GDF15* expression. (**G**) Boxplots of *ISR^GDF15^* scores for each condition, sorted by ascending median score. P-values from Kruskal-Wallis test and Dunn’s test for multiple comparisons. *Oligo,* Oligomycin; DEX = dexamethasone, mitoNUITs = mitochondrial nutrient uptake inhibitors, 2DG = 2-Deoxy-D-glucose, BHB = β-hydroxybutyrate, contact inhib = contact inhibition.

We used a longitudinal transcriptomic dataset including 339 fibroblast samples collected from either healthy donors or mitochondrial-disease donors with mutations in the OxPhos complex IV assembly-factor gene *SURF1* (**Figure 1B**). To create a vast spectrum of bioenergetic conditions, each cell line was exposed to up to 11 metabolic perturbations including an inhibitor of the OxPhos system, oligomycin (Oligo), glucocorticoid agonist dexamethasone (DEX), mitochondrial-nutrient-uptake inhibitors (mitoNUITs), glucose deprivation plus β-hydroxybutyrate (BHB), low oxygen (3%), and contact inhibition^13^. OxPhos defects are potent triggers of the ISR in other datasets^1^ and in this dataset^14^, making this model and the variety of conditions tested an ideal testbed to examine gene expression signatures of the GDF15 arm, and other dimensions of the ISR, in human cells (**Figure 1C**).

To computationally isolate co-regulated ISR genes, we queried bulk gene expression RNAseq data using *GDF15* as a bait; we included *GDF15* alongside the 119 ISR genes from the three ISR gene lists and performed dimensionality-reduction analyses (the 120 genes can be found in **Extended Data Table 1**). This allowed us to explore gene expression patterns within our samples, with a focus on patterns most strongly associated with *GDF15*. Using principal component analysis (PCA) to explore this gene-expression landscape across all treatments simultaneously (**Figure 1D**), we identified a major axis of genes co-regulated with *GDF15* (principal component 2, explains 15.5% of variance). **Extended Data Figure 2A** shows the loadings for each gene on each of the first two principal components.

Next, we used factor analysis to more quantitatively isolate the transcriptional signature of this *GDF15*-related ISR axis. This approach converged on an optimal solution containing 12 factors (i.e., sets of loadings or “weights” applied to each of the 119 genes plus *GDF15* for a total of 120 genes), which collectively account for a maximal portion (73.1%) of the shared variance within this dataset (**Figure 1E**). The relationships between the principal components and each factor are quantified in **Extended Data Figure 5A**, with specific correlation plots depicted in **Extended Data Figure 5B-C**. Factor 1—our index hereafter referred to as the *ISR^GDF15^* score—explained 12.9% of the variance, and was more strongly correlated with *GDF15* expression (Spearman’s rank correlation coefficient (ρ) = 0.87, *p*<1×10^-15^) than any other factor, establishing construct validity for the calculated *ISR^GDF15^* score (**Figure 1F**). The 11 remaining factors and their gene loadings, which are not explored in this work but available to readers in **Extended Data Table 2**, may represent additional pathways indicative of other ISR dimensions.

To understand the biology underlying the *ISR^GDF15^* index, we compared the genes most strongly associated (positively or negatively) with *GDF15*. The genes contributing most strongly to the *ISR^GDF15^* index were positively enriched for regulation of endoplasmic reticulum–unfolded protein response [top 10_positive_: *DDIT3*, *PPP1R15A*, *ERN1*, *GADD45A*, *CBX4*, *BBC3*, *SLC3A2*, *SIAH2*, *CEBPB*, *TRIB3*]. In contrast, negatively weighted genes for the *ISR^GDF15^* index were enriched for the regulation of translation and translation initiation [top 10_negative_: *PPP1CA*, *APAF1*, *SLC35A4*, *FOS*, *IMPACT*, *NAIP*, *BIRC5*, *EIF2B1*, *NARS1*, *GCN2*] (see **Extended Data Table 3** annotations and known functions of each gene), a well-described consequence of ISR activation in most cell types. These results are also consistent with the notion that cellular processes operate under energy constraints: that is, while the activation of some energy-dependent processes are upregulated, other processes must be downregulated^15,16^. As such, cells activating the GDF15-signaling ISR arm may generally downregulate protein synthesis, as previously observed^17^, despite known increases in specific proteins, including GDF15^14,18^.

Functionally, *ISR^GDF15^* was most significantly induced in oligomycin-treated cells (large effect size, *g* = 2.13, *p*< 0.001) and in SURF1-mutant cells (*g* = 1.91, *p*<0.001, **Figure 1G**) relative to untreated control cells, as expected. The conditions of no glucose with β-hydroxybutyrate (*g* = 1.33, *p*<0.001) and the glycolysis inhibitor 2-deoxyglucose (*g* = 0.97, *p*<0.05) also resulted in a higher *ISR^GDF15^*, which is consistent with glucose deprivation being a known ISR activator^19^. The higher *ISR^GDF15^* observed in the contact-inhibition condition (*g* = 1.23, *p*<0.001) was surprising at first, as it was expected to be more physiologically similar to what cells experience in vivo (i.e., a more physiological skin cell culture). However, the culture protocol used in this condition called for only one media change per week (matching the passaging of the non-contact-inhibited cells). This protocol led to a higher density of cells throughout the week, which, in turn, may have depleted nutrients at a faster rate. Thus, the ISR was likely activated in the contact-inhibition group as a result of nutrient starvation.

To our initial surprise, the low oxygen level (3%, not anoxia) treatment was not an activator of the ISR, as would be logically expected of a hypoxic treatment. However, upon reviewing the literature, we noted that the experimental conditions previously considered hypoxic treatments for these fibroblast cells was, in fact, likely a physiological oxygenation level^20^. In vivo skin has oxygenation levels between 1.1-4.6%^20^, and thus is likely to be a better control than our control group, which is cultured at 21% oxygen. Together, these data demonstrate the existence of a relatively small subset of positively and negatively inducible *GDF15*-related ISR genes induced by mitochondrial OxPhos perturbations.

To validate the *ISR^GDF15^* index, we then tested how targeted and time-resolved ISR activation influenced GDF15 itself, and the *ISR^GDF15^* index. To achieve this, we used data from a light-activatable protein kinase R (PKR) kinase in the H4 cancer line (**Figure 2A**)^21^. This optogenetic model triggers robust activation of the ISR (marked by p-eIF2a, ATF4, and CHOP induction) without other confounds including the simultaneous activation of other pathways, such as damage response pathways, for example.

**Figure 2.**
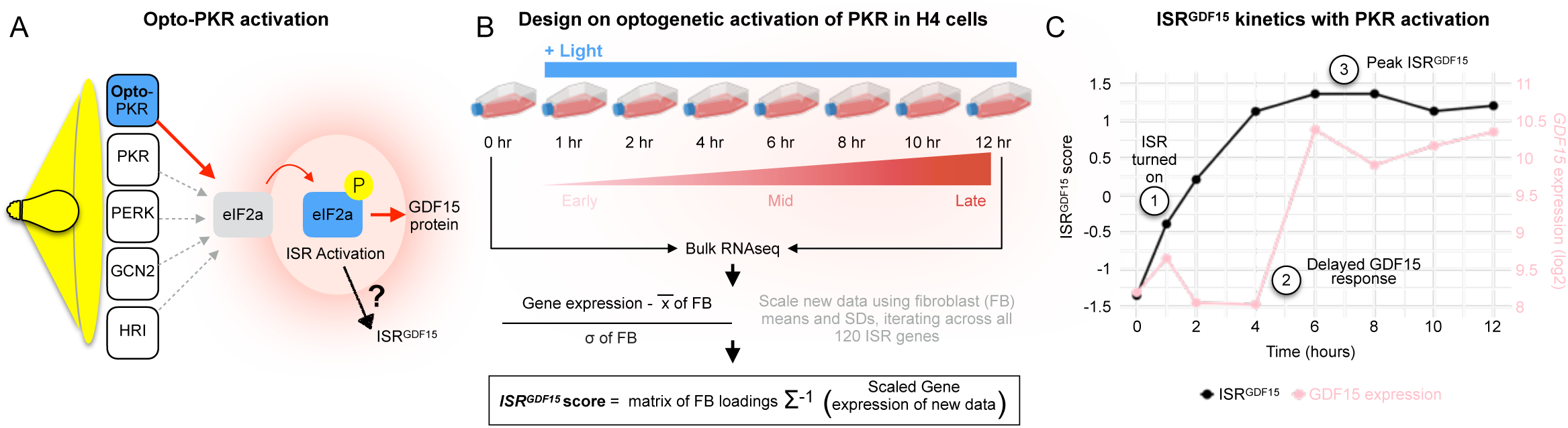
Validation of the ISR^GDF15^ index with time-resolved ISR activation. (**A**) Optogenetic ISR activation strategy targeting an engineered light-sensitive PKR, an ISR kinase, which phosphorylates eIF2alpha to activate the ISR, as per Batjargal et al. (**B**) Experimental and data processing scheme to compute the *ISR^GDF15^* scores across 8 timepoints spanning 12 hours of continuous PKR activation in H4 cells. (**C**) *ISR^GDF15^* score and *GDF15* RNA transcript levels for the H4 cells under increasing hours of ISR stimulation.

The light sensitive–PKR H4 cells were stimulated with light for up to 12 hours, and cells harvested at eight timepoints for bulk RNAseq (**Figure 2B**). We deployed the *ISR^GDF15^* index to the RNAseq data, quantifying its expression at each timepoint (See **Figure 2C**). The *ISR^GDF15^* score increased by 1 hour and peaked at 6-8 hours, followed by a plateau over the remaining hours of light stimulation. The trajectories of induction for other factor scores are shown in **Extended Data Figure 6A**.

Comparatively, PKR activation did not induce *GDF15* expression over the first 4 hours. However, between hours 4 and 6 we observed a ∼5-fold increase in GDF15 mRNA at 6 hours (**Figure 2C**). Together, these findings support the accuracy and sensitivity of the *ISR^GDF15^* index to PKR-mediated ISR activation, even in the absence of changes in *GDF15* expression from 0 to 4 hours. This suggests that the *ISR^GDF15^* index captures ISR biology more broadly and sensitively than *GDF15* alone.

To examine the extent to which the *ISR^GDF15^* index it is conserved in human tissues, we used the Genome-Tissue Expression (GTEx) project, which includes RNAseq data on up to 17 different organs from ∼1,000 donors^22^ (**Figure 3A left**). We first asked whether the *ISR^GDF15^* score was correlated with its bait *GDF15* gene, when applied to individual human organs and tissues. If the internal correlation structure and loadings of the genes composing the *ISR^GDF15^* index were similar across datasets, we expected positive correlations between the index and the gene *GDF15*. Focusing on tissues with a sample size >20, we observed significant positive correlations between the *ISR^GDF15^* and *GDF15* in 43 out of 44 (97.7%) human tissues (ρ = 0.41-0.82, all *p*’s <0.05 corrected for multiple testing, **Figure 3B**). These findings suggest that the genetic architecture of the *GDF15*-related arm of the ISR is at least partially conserved between fibroblasts in vitro and human tissues in vivo.

**Figure 3.**
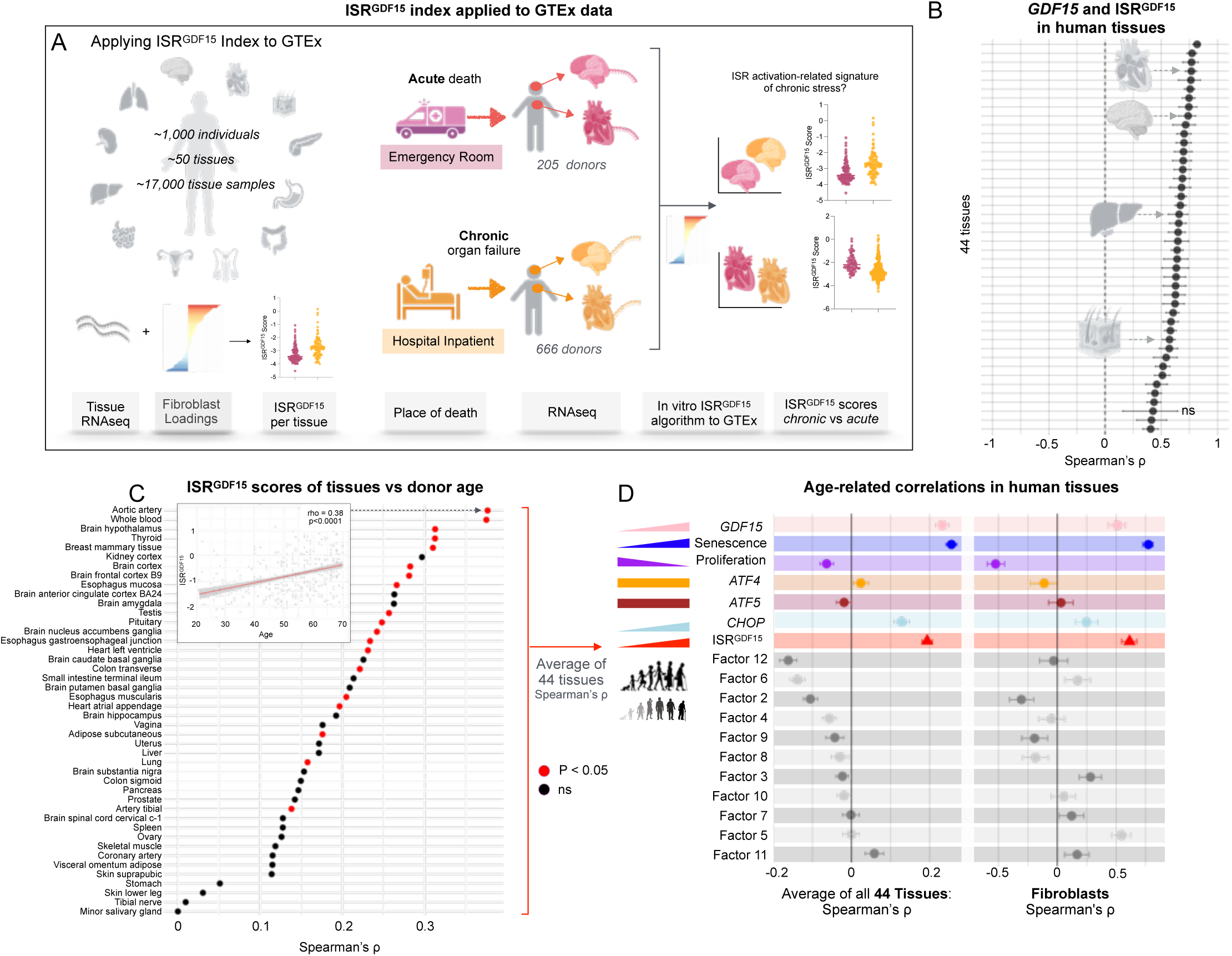
ISR^GDF15^ index applied to human GTEx dataset correlated with age at death. (**A**) Schematic of human GTEx dataset containing a variety of human organ samples, and applying the fibroblast generated index to the human transcriptomic data to get the human *ISR^GDF15^* scores for each tissue. (**B**) *ISR^GDF15^* correlations with *GDF15* across all the tissue samples expressing all 120 genes in the *ISR^GDF15^* index (Spearman’s ρ, +/- 95% confidence intervals, Bonferroni corrected for multiple comparisons). (**C**) Spearman’s rank correlations of each tissue’s *ISR^GDF15^* score compared to donor age (red points denoting a Bonferroni corrected p<0.05). (**D**) Average Spearman’s correlations (+/- SEM) across all 44 tissues comparing the donor age with the chosen gene(s) or *ISR^GDF15^* score or factor scores (left) and fibroblasts’ age correlated with the chosen genes or *ISR^GDF15^* score or factor score (+/- 95% confidence interval) (right). *ISR^GDF15^* average tissue correlation denoted with a red triangle. Wilcoxon signed-rank test used compared against 0.

Given the well-known upregulation of GDF15 with age^2^, but not necessarily of the ISR per se, we examined the association of *ISR^GDF15^* score with age in GTEx (range 20-70, average 53 years). The correlations were positive as expected in all human tissues, and 33 of 44 tissues were significant (average ρ = 0.23, *p*<0.001, **Figure 3D**). Seeking a generalizable test to examine how different gene expression-based indices perform across fibroblasts and human tissues, we average expression across all tissues and individuals of different health status included in the dataset. GDF15 alone also was correlated with age, and in some tissues more strongly so than the *ISR^GDF15^* score (**Figure 3D**).

To pressure test this analysis further, we confirmed that senescence-associated genes (averaged tissue expressions of *CDKN1A*, *CDKN2A*, *CCND2*) were upregulated, whereas proliferative or replication-related genes (averaged tissue expressions of *TOP2A*, *RRM2*, *MKI67*) were downregulated with advancing age, validating the use of cellular lifespan system^13^ to probe time- or age-related changes. Other single ISR genes *ATF4* and *ATF5* were unrelated to age, whereas *CHOP* was mildly related to age (ρ = 0.13, *p*<0.001) with about half the effect size of *GDF15*. In sum, *ISR^GDF15^* was positively associated with age (ρ = 0.19, *p*<0.001), indicating that the *GDF15*-related arm of the ISR is upregulated with age, and more strongly than individual genes classically referenced alone (or in conjunction with others) to denote activation of the ISR (**Figure 3D**).

We then systematically examined the other 11 potential arms of the ISR isolated by factor analysis, ranked from negative to positive age-related effect sizes across all GTEx tissues (**Figure 3D**). Four other ISR arms were significantly negatively associated with age (Factors 12, 6, 2, 4), and one was positively associated with age (Factor 11). Applying the same analysis to aging human fibroblasts similarly showed both *GDF15* alone (ρ = 0.51, *p*<0.001) and the *ISR^GDF15^* score (ρ = 0.61, *p*<0.001) were positively associated with age, though these effect sizes were smaller than the senescence associated markers (ρ = 0.77, *p*<0.001), as expected.

A direct comparison of effect sizes across each gene, the *ISR^GDF15^ score*, and the 11 factors revealed a moderate correlation between the in vivo and in vitro datasets (ρ = 0.69, *p*<0.01, **Extended Data Figure 7A left**). The correlation was stronger without the 11 non-*ISR^GDF15^* factors (ρ = 0.93, *p*<0.01, **Extended Data Figure 7A right**). Given the coherence between the two systems (in vivo and in vitro), this finding supports the notion that the *ISR^GDF15^* signaling arm has an age-related association similar to that of *GDF15* in both strength and direction. In other words, the *ISR^GDF15^* index accounts for most of the age-related signal of the ISR. The other factors may be important processes, but their age associations are considerably weaker than the identified *ISR^GDF15^* arm.

In the GTEx dataset, some donors died suddenly in the intensive care unit (76.4% of a cardiac arrest or related cause), while other individuals were hospital inpatients that endured more protracted death processes (involving coma, or other conditions). To test whether ISR activation was associated with organ-specific cause of death, we calculated the *ISR^GDF15^* score for each tissue for each place of death: emergency room (ER) and hospital inpatient (HI). We focused on these groups for two reasons: i) each group included a sufficient number of samples to allow robust comparisons of ISR activation; and ii) acute ER deaths are more likely to have been associated with acute cardiocirculatory death within hours, while HI deaths are more likely to have endured metabolic disturbances and an protracted agonal period that tends to affect brain function (e.g., coma). Because the ISR is activated in a time-dependent manner^7^, we hypothesized that longer hospital stays and pathology in HI would produce higher *ISR^GDF15^* activation compared to ER donors, who on average had less time to activate the ISR before death.

The expected ISR activation pattern was only observed in a specific subset of tissues: brain, salivary gland, skeletal muscle, skin, tibial nerve and artery, and subcutaneous adipose tissue (no digestive tissues). We also found that donors from the ER had significantly higher *ISR^GDF15^* scores in a different subset of tissues: whole blood, mammary breast, prostate, thyroid, esophageal, lung, transverse colon, stomach and the atrial appendage, left ventricle, aorta, and coronary artery of the heart; not in CNS tissues (**Figure 4C**).

**Figure 4.**
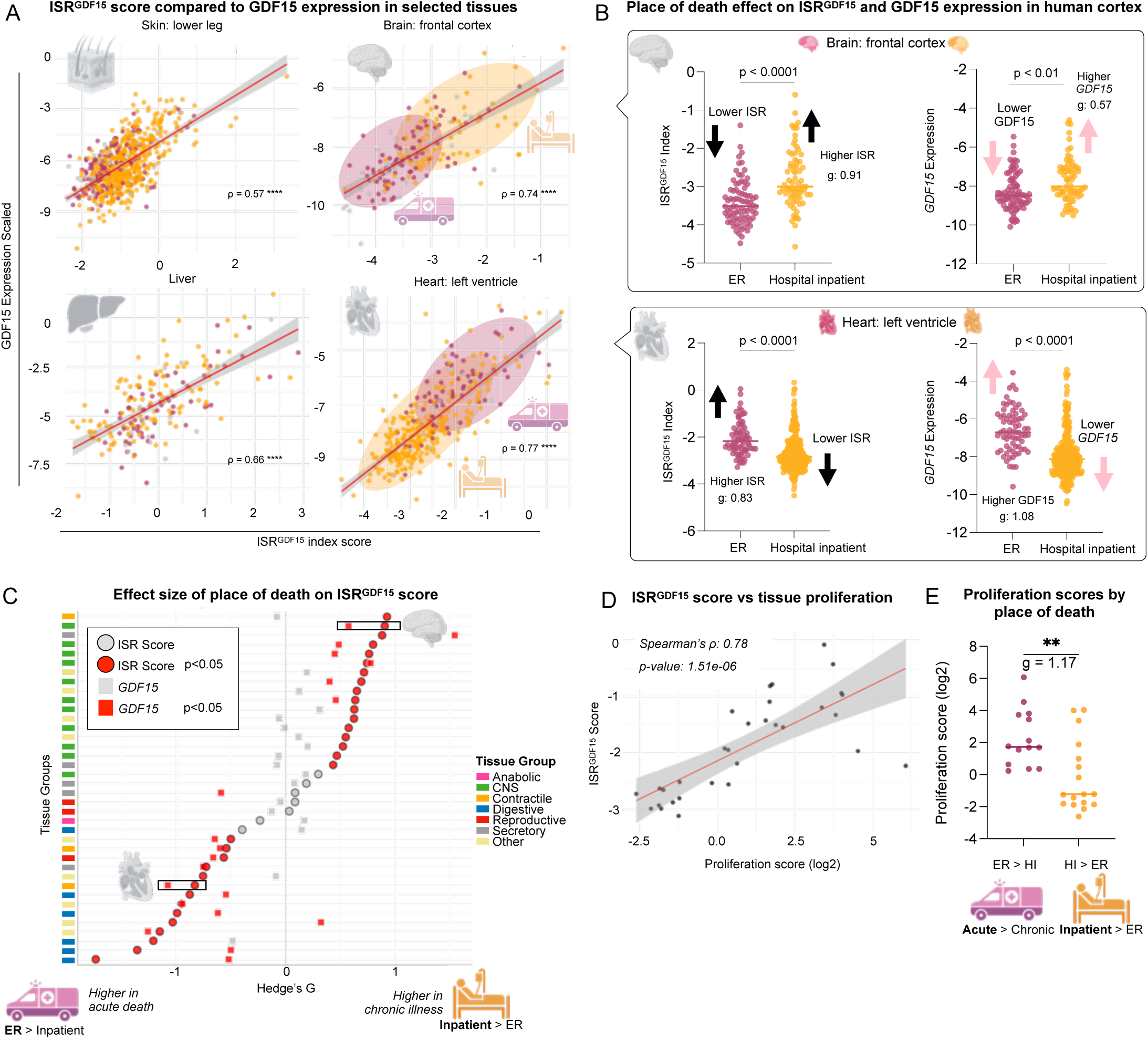
*ISR^GDF15^* index applied to human GTEx dataset. (**A**) Select tissue correlations of *ISR^GDF15^* and *GDF15* expression, Bonferroni corrected. (**B**) *ISR^GDF15^* or *GDF15* expression for each place of death, in brain frontal cortex or heart left ventricle; significance determined via Wilcoxon rank-sum test. (**C**) Hedge’s g effect sizes comparing each tissue’s *ISR^GDF15^* activation score for each place of death, or *GDF15* expression comparing each place of death—Hedge’s g values of hospital inpatient scores being greater than ER seen on the right, ER greater than hospital inpatient towards the left. Circles denote *ISR^GDF15^* score, squares *GDF15* expression, red as significant, gray non-significant, with BH correction for multiple comparisons. The grey brain and heart indicate the *ISR^GDF15^* score and *GDF15* expression of the brain frontal cortex and the heart left ventricle comparisons from **B**. See methods for specific tissues within each tissue grouping. Wilcoxon rank-sum test was used to compare either the *ISR^GDF15^* score or *GDF15* expression in ER or HI, with BH correction. (**D**) Proliferation scores of each tissue split into two groups based on whether the tissue had a higher ISR score in ER or HI from Figure 2C. (**E**) Spearman’s correlation of *ISR^GDF15^* scores with proliferation scores. ER = Emergency Room, HI = hospital inpatient.

To investigate this unexpected pattern, we drew upon previous research by Mick et al., which demonstrated that the same stressor triggered differential ISR activation between myoblasts (actively proliferating) compared to myotubes (non-proliferative)^1^. To explore whether proliferation status could contribute to the differential ISR activation across human tissues, we calculated the proliferation score of each tissue ^23^ (see methods for details) and split the tissues into two groups: proliferative vs non-proliferative, using the median proliferation score. We then compared the *ISR^GDF15^* scores between the two groups based on their proliferation status. Proliferative tissues exhibited significantly higher ISR activation scores than the non-proliferative tissues (**Extended Data Figure 8A**). This was validated by the significant correlation between ISR activation scores and proliferation scores across tissues (ρ = 0.78, *p*<0.001, **Figure 4D**). Finally, we categorized the tissues according to whether the donors had significantly higher *ISR^GDF15^* scores based on deathplace (either HI to ER or ER to HI), and found that the two groups had different proliferative indexes (*g* = 1.17, *p*<0.01, **Figure 4E**). The tissues with higher *ISR^GDF15^* in the ER group were more proliferative than those that were higher in the HI group. Ultimately, these analyses support the notion that proliferation status contributes to differential ISR activation in a tissue-dependent manner, and help explain the distribution of ISR activation observed across tissues according to place of death seen in **Figure 4C**.

Interestingly, in these post-mortem tissues, *ISR^GDF15^* was more sensitive than *GDF15* expression to sudden death vs chronic illness (or vice versa), as reflected by 81.6% of tissues showing significantly different *ISR^GDF15^* scores under these two scenarios compared to only 52.6% of tissues showing a significant difference in *GDF15* expression under the same scenarios (Chi square = 5.96, *p* = 0.015, **Figure 4C**). This was exemplified in the frontal cortex of the brain, which displayed a large effect size (*g* = 0.91) for *ISR^GDF15^* when comparing sudden death to chronic illness, but only a moderate effect size for *GDF15* (*g* = 0.57, **Figure 4B**). These results are meaningful, particularly in the context of a robust correlation between *ISR^GDF15^* and *GDF15* expression in this tissue (ρ = 0.74, **Figure 4A**). This supports the use of the *ISR^GDF15^* score rather than *GDF15* alone as an indicator of ISR activation.

## Discussion

Together, this work challenges the monolithic understanding of the ISR in humans, showing instead that the ISR—as defined by popular gene lists^10–12^—contains distinct sub-responses or sub-pathways. In particular, our results highlight potential tissue-specificity in ISR activation with disease and conditions leading up to death (see **Figure 4C-E**). These findings call for the consideration of tissue-specific regulatory mechanisms or downstream effects that have yet to be defined. We also identify a conserved set of *GDF15*-related canonical ISR genes upregulated by experimental mitochondrial perturbations, as well as other groups of genes (factors 2-12) that reflect the multidimensional nature of the ISR. The identified *ISR^GDF15^* index and *GDF15* expression were co-regulated across most human tissue, and some aspects of the ISR genetic architecture (*ISR^GDF15^* in particular) were found to be conserved across three distinct datasets, increasing our confidence in the robustness of these findings. Finally, this work illustrates that *ISR^GDF15^* is activated in a time-dependent manner that precedes *GDF15* expression, making it an indicator of early (pre-GDF15 induction) ISR detection.

The limitations of this study include i) the use of linear statistical methods, while genes likely interact in non-linear ways^24^, ii) analyses from post-mortem human tissues, where the post-mortem period could alter gene expression profiles, at least in the brain^25^, iii) the optogenetic experiments were only performed with a single ISR kinase, which, while sufficient to robustly induce the ISR^21^ may not paint the full picture of ISR biology. We also note that the genes from the literature-based consensus lists could be missing other important genes. Further work to demonstrate the activation or inactivation of the index could be performed with the use of ISR inhibitors such as ISRIB, which would provide a negative control that was not available in these experiments. This would be especially useful for opto-PKR, given the relatively high baseline expression of *GDF15* observed here, which is consistent with high GDF15 expression in several cancer cell lines (see Human Protein Atlas^26^).

Overall, we present a general approach to develop multi-gene indices, and the *ISR^GDF15^* index in particular provide a foundation for future research aiming to dissect the multifaceted biology of the human ISR in relation to health, disease, and aging.

## Methods

### ISR Gene List Compilation

Gene lists for the ISR were gathered by inputting the phrase “integrated stress response” into the search bars of the main websites for three sources: GO Consortium, The Jackson Laboratory, and Any Genes^10–12^. The genes from each lists were merged to make one combined ISR gene list. Some gene names were altered to fit the nomenclature of the specific transcriptomic dataset being used, as different gene aliases were used in each dataset for some genes (i.e., NARS1/NARS, WARS1/WARS, Igtp/IRGM). The genes found in each gene list are reported in **Extended Data Table 1**. Information about the top 10 positively and negatively associated genes in the *ISR^GDF15^* index was found by searching each gene in URL: https://www.genecards.org/.

### Fibroblast Lifespan Study

The lifespan study is explained in full detail in Sturm et al^13^. RNAseq data was downloaded from the shiny app URL: https://columbia-picard.shinyapps.io/shinyapp-Lifespan_Study/. Fibroblast transcriptomic data was previously normalized via DEseq2, v1.30.1, and was log2 transformed. A total of 339 samples (excluding control line technical replicate data) was used for further analysis. Five healthy control lines and three patient-derived SURF1 mutant lines were cultured, with or without various metabolic treatments, as described in Sturm et al.^13,14^.

### PCA

R version 4.4.1 with R studio was used to perform PCA (and all subsequent statistical analyses). The ‘prcomp’ function was used to generate PCA, (*stats* package v4.4.1) using 339 fibroblast samples and the master list of ISR genes plus *GDF15*. Principal component biplots of components beyond PC1 vs PC2 are plotted in **Extended Data Figure 3**, as well as their correlations against fibroblast days in culture in **Extended Data Figure 2B-C**.

### Factor Analysis

The same fibroblast dataset of 339 samples and 120 genes was used in the factor analysis. The ‘factanal’ function (*stats* package) was used, with a varimax rotation. The scree method^27^ was used to determine the number of factors, as no predetermined number was hypothesized given that the number of causes and outputs of the ISR have yet to be determined. Of the 119 ISR genes, the *ISR^GDF15^* index included 77 positive and 42 negative gene loadings, reflecting ISR genes positively and negatively related to *GDF15* (see **Extended Data Table 4** for individual gene loadings). Individual genes, such as *GDF15*, were centered and scaled by using the scale() function in R before comparing against the *ISR^GDF15^* score or age. A biplot of *ISR^GDF15^* (Factor 1) vs Factor 2, as well as all factor scores compared to age are depicted in **Extended Data Figure 4**.

### Opto-PKR ISR Score

Raw FPKM values were normalized to TPM from Batjargal et al. and was log2 transformed. Then, the values were centered and scaled to the initial fibroblast dataset, and finally the *ISR^GDF15^* index was applied, as depicted in the bottom half of **Figure 2B**.

### GTEx

RNA sequencing data was downloaded as gene expression read counts from GTEx Analysis V8. RNA integrity number (RIN) values of less than 6 were filtered out of the data. Trimmed Mean of M-values (TMM) values were calculated using DGE method (BiocGenerics package, version 0.50.0). To convert the human TMM values to the *ISR^GDF15^* score, human data were scaled to the initial fibroblast dataset, as depicted in **Figure 2B** (by subtracting the human TMM values by fibroblast means and dividing by the fibroblast standard deviation), and applied to the correlation matrix of the initial fibroblast data to the human data. Tissue samples of less than 20 were excluded from the analysis, leaving the kidney cortex with the minimum number of tissue samples of 86. Four tissues were excluded from the application of the *ISR^GDF15^* index due to missing one or more genes from the ISR list: bladder, brain cerebellar hemisphere, brain cerebellum, and adrenal gland. A total of 44 tissues were included in the analyses. Three genes required changing aliases to match the fibroblast genes: *DELE1* was changed to *KIAA0141*, *WARS1* was changed to *WARS*, and *NARS1* was changed to *NARS*. When comparing either the *ISR^GDF15^* score of ER vs HI or *GDF15* of ER vs HI, tissues with less than 20 samples per group (ER or HI), were filtered out. Six tissues were filtered out: kidney cortex, pancreas, small intestines of the ileum, spleen, uterus, and vagina for having less than 20 samples in the ER group. For distinguishing cardiac-related deaths within the HI or ER groups, we included the following terms as cardiac-related under the cause of death columns either DTHCOD or DTHFUCOD: “cardiac”, “acute mi”, “myocardial”, “myocardiac”, “arrest”, “cardio”, “bradycardia”, “heart”, “aortic dissection”, “probable mi”, and “chf”. Causes of death that did not include any of those terms were deemed non-cardiac. The HI group was composed of 25.2% cardiac-related deaths. Various tissues in the GTEx dataset were noted in the pathology notes column to contain other tissues (often fat) these samples were not filtered out.

### Tissue Groups

As described in Devine et al.^23^, tissues were split into the following groups: liver into Anabolic; all brain tissues into central nervous system (CNS); skeletal muscle, heart atrial appendage and heart left ventricle into Contractile; colon tissues, esophageal tissues, and stomach into Digestive; testis, ovary, and prostate into Reproductive; salivary gland, adipose subcutaneous and visceral omentum, thyroid, and pituitary into Secretory; skin lower leg and suprapubic, artery tibial, breast mammary, whole blood, artery coronary, lung, and artery aorta into Other.

### Senescence and Proliferation Index Scores

Genes for senescence (*CDKN1A*, *CDKN2A*, *CCND2*)^28^ or proliferation (*TOP2*, *RRM2*, *MKI67*)^23^ were selected to create either a tissue senescence score or proliferation score, by averaging the three senescent or proliferative genes respectively. Selection criteria for the proliferation index genes are explained in the methods of Devine et al.^23^. When splitting the tissues into two groups, proliferative vs non-proliferative, the median score of 0.63 was used.

### Statistics

All correlations performed were Spearman’s rank correlations. When comparing group differences of two groups, Wilcoxon rank-sum tests were used. When noted, Bonferroni correction for multiple comparisons was used, with a p-value threshold of 0.05. Otherwise, we used the Benjamini-Hochberg (BH) method, a less stringent form of multiple testing corrections, for more exploratory questions. Effect sizes were determined via Hedge’s g, an unbiased analog of Cohen’s d^29^. Hedge’s g was calculated via the R function ‘cohen.d’ from the package *effsize*, with the Hedge’s g correction applied. When comparing multiple independent treatment groups simultaneously, Prism version 10.2.3 (347) was used to perform a Kruskal-Wallis test followed by Dunn’s test for multiple comparisons. Wilcoxon signed-rank test was performed using Prism, when testing if a group significantly differed from 0.

## Acknowledgements

Some figure elements were generated with Biorender.com. We are grateful to the GTEx consortium and to Gabriel Sturm for creating the valuable Cellular Lifespan Study data resources that made this work possible.

## Author contributions

J.L.M.S and M.P. conceived the project. J.L.M.S. performed the analyses with K.T. J.L.M.S. generated the figures. J.L.M.S., M.P., A.A.C., K.T., A.S.M., and J.D. interpreted the data. T.B. and M.Z.W provided data for the optogenetic ISR activation. J.L.M.S., and M.P. drafted the manuscript with A.A.C. All authors reviewed and edited the final manuscript.

## Data and code availability

GTEx v8 RNAseq data is available for download at https://gtexportal.org/home/downloads/adult-gtex/bulk_tissue_expression.Cellular. The Lifespan Study fibroblast dataset can be downloaded at https://columbia-picard.shinyapps.io/shinyapp-Lifespan_Study/. R code for the analyses will be made available at www.github.com/mitopsychobio.

## Extended Data Material

This article contains 8 Extended Data Figures and 4 Extended Data Tables.

---

**Extended Data Figure 1.**
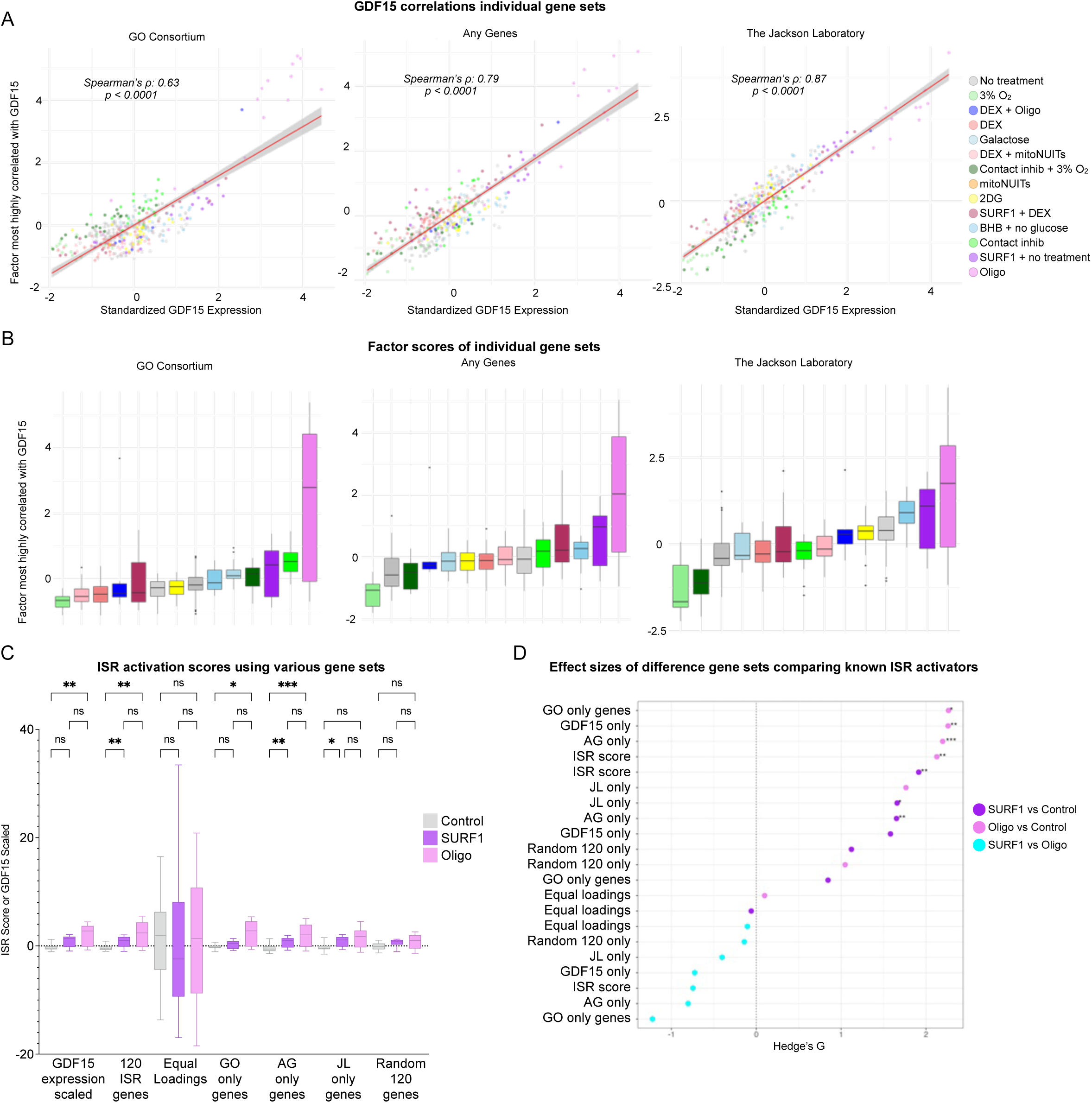
Sensitivity analysis for various ISR gene lists. (**A**) Factor analysis performed on individual gene sets (GO Consortium which contained 37 genes, Any Genes which had 84 genes, and Jackson Laboratory which had 23 genes), as well as equal loadings for all the 120 genes (not shown), and the average of 20 different sets of 119 randomly chosen genes plus GDF15 (not shown). The factor that was most correlated with *GDF15* was chosen for further sensitivity analyses. (**B**) The index scores of the chosen factors from individual gene sets depicted in **A**. (**C**) Comparison of the index scores of only control, SURF1 mutant, or Oligo treated cells for each gene set, including equal loadings of the 120 ISR genes; Kruskal-Wallis test performed with Dunn’s test for multiple testing. (**D**) The effect sizes of each treatments comparisons (Hedge’s g) from panel **C**. GO = GO Consortium, AG = Any Genes, JL = Jackson Laboratory

**Extended Data Figure 2.**
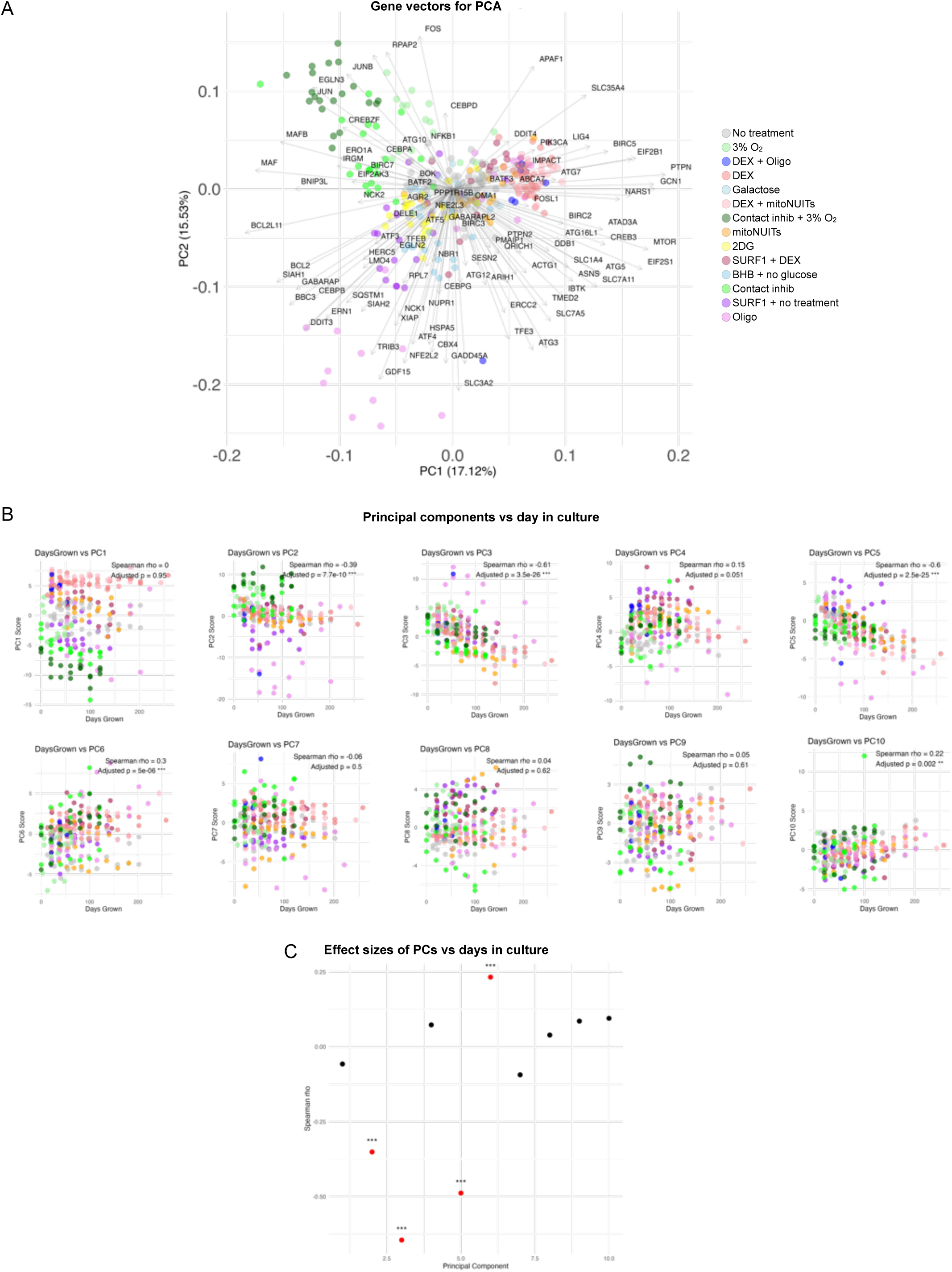
Principal component gene vectors and comparisons with age. (**A**) Biplot with gene vector loadings of PC1 and PC2. (**B**) First 10 PC components vs fibroblast time in culture (days). (**C**) Effect sizes (Spearman’s rank correlations) of PCs vs time in culture (days).

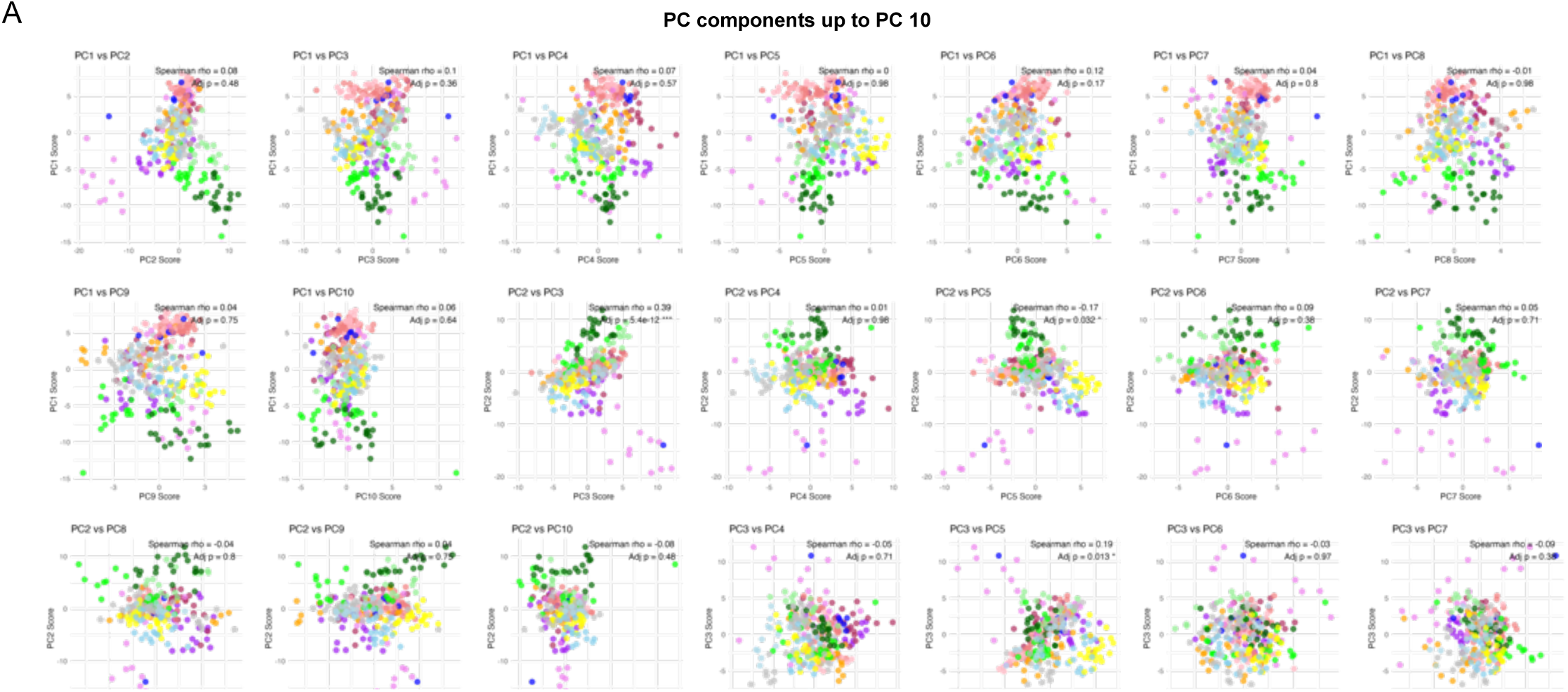

**Extended Data Figure 4.**
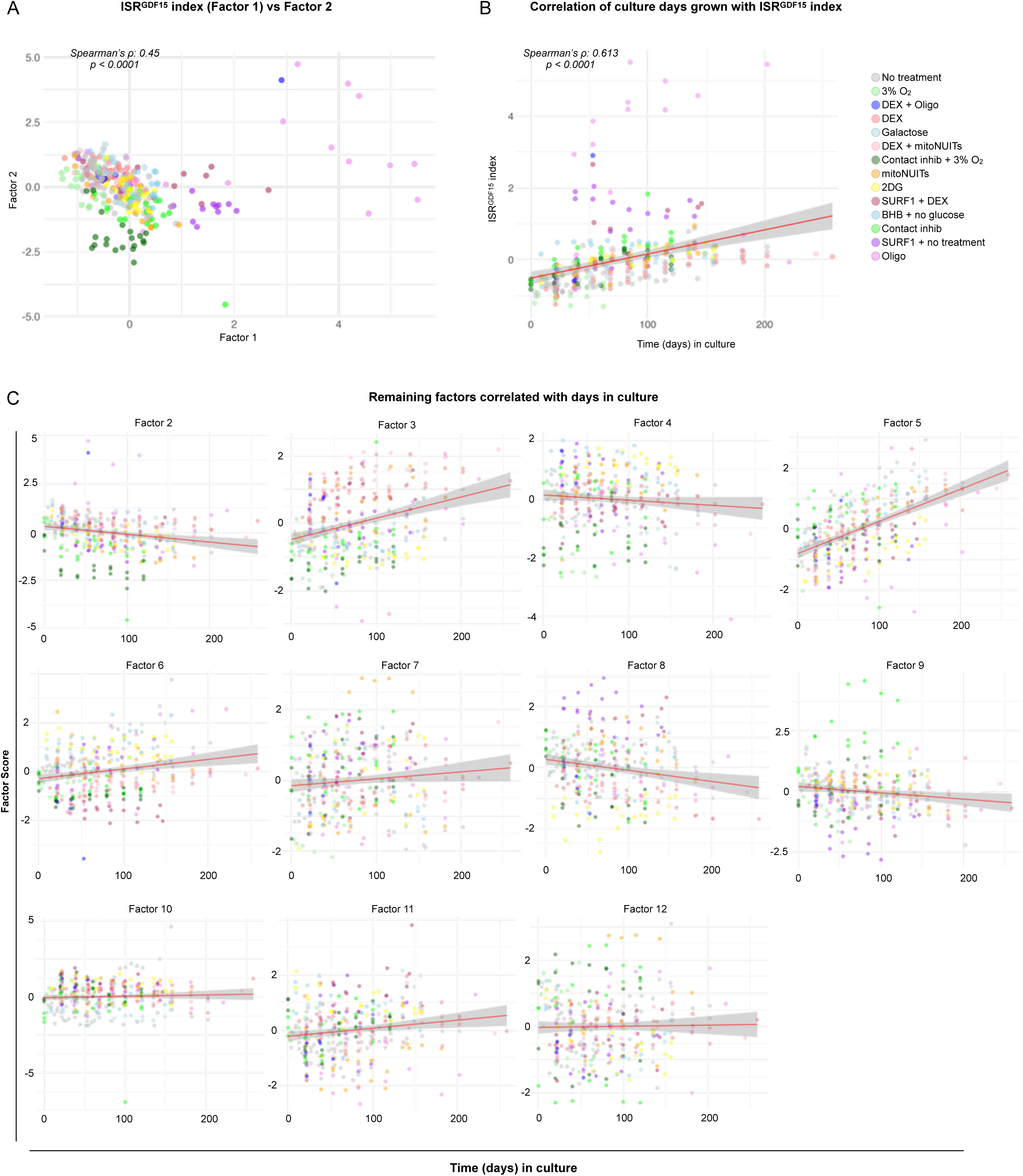
Factor analysis comparisons with age. (**A**) Plot of first two factors from the factor analysis. (**B**) Correlation of each fibroblast sample vs time (in days) of culture. (**C**) Remaining 11 factors scores vs time in culture.

**Extended Data Figure 5.**
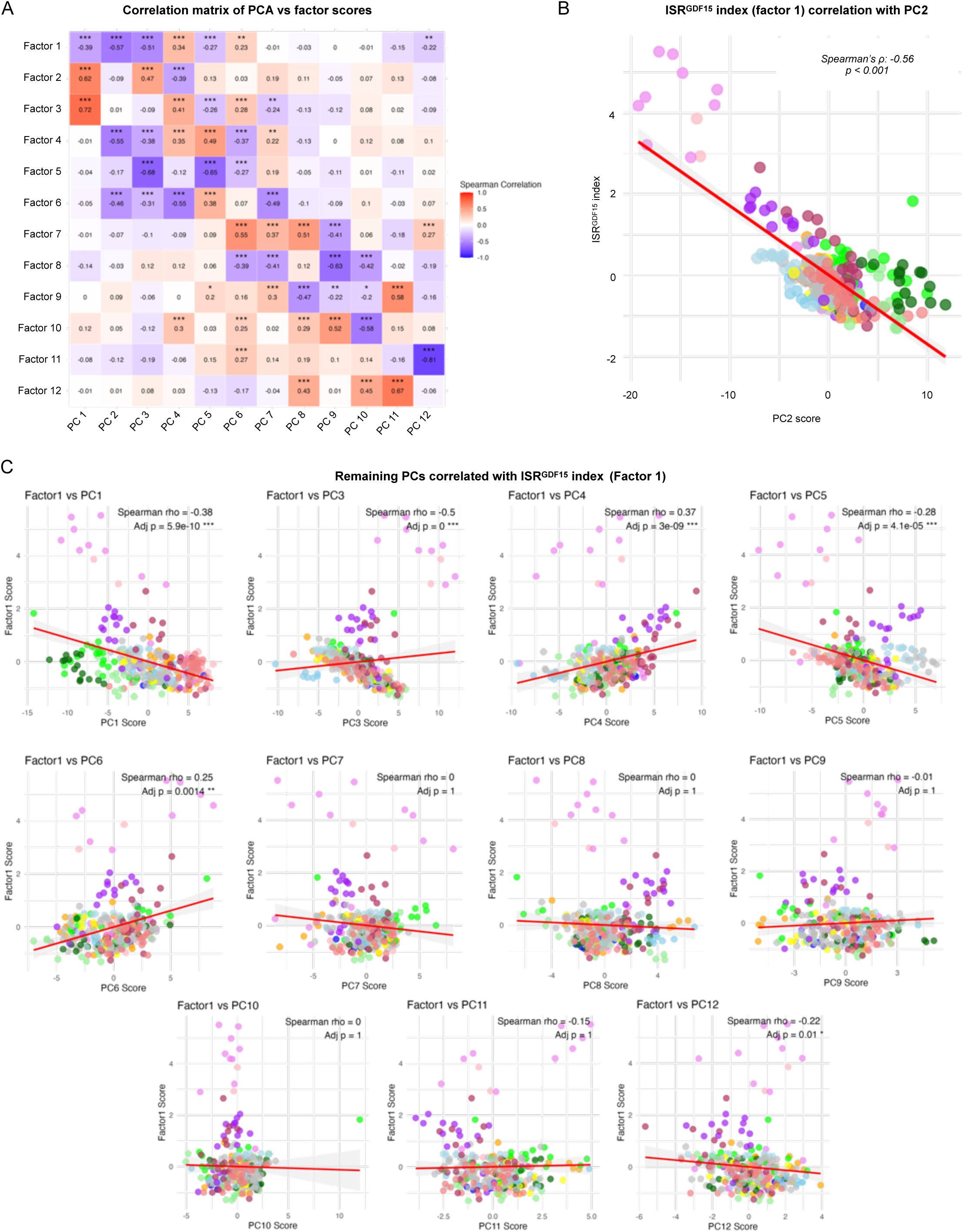
Correlations of factor scores and principal components. (**A**) Correlation matrix of Spearman’s rank correlations of the factor scores vs the first 12 principal components from the PCA. (**B**) Spearman’s correlation of *ISRGDF15* index (Factor 1) vs PC2 score. (**C**) Spearman’s correlations of *ISRGDF15* vs remaining principal components up to 12.

**Extended Data Figure 6.**
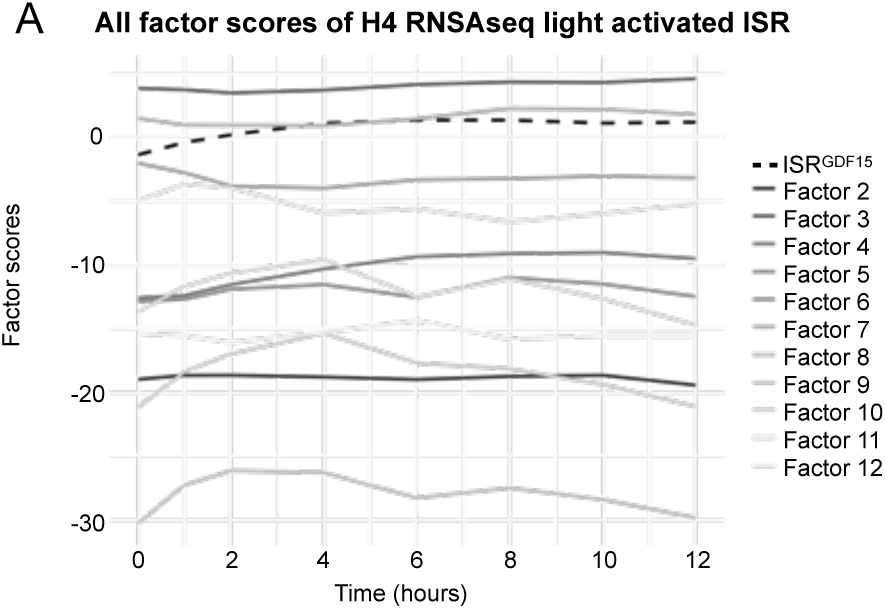
All factors applied to opto-PKR H4 cells. (**A**) Each factor score for the H4 cells, containing the ISR sensor kinase PKR engineered to be activate in response to light (opto-PKR), stimulated with 0–12 hours of light.

**Extended Data Figure 7.**
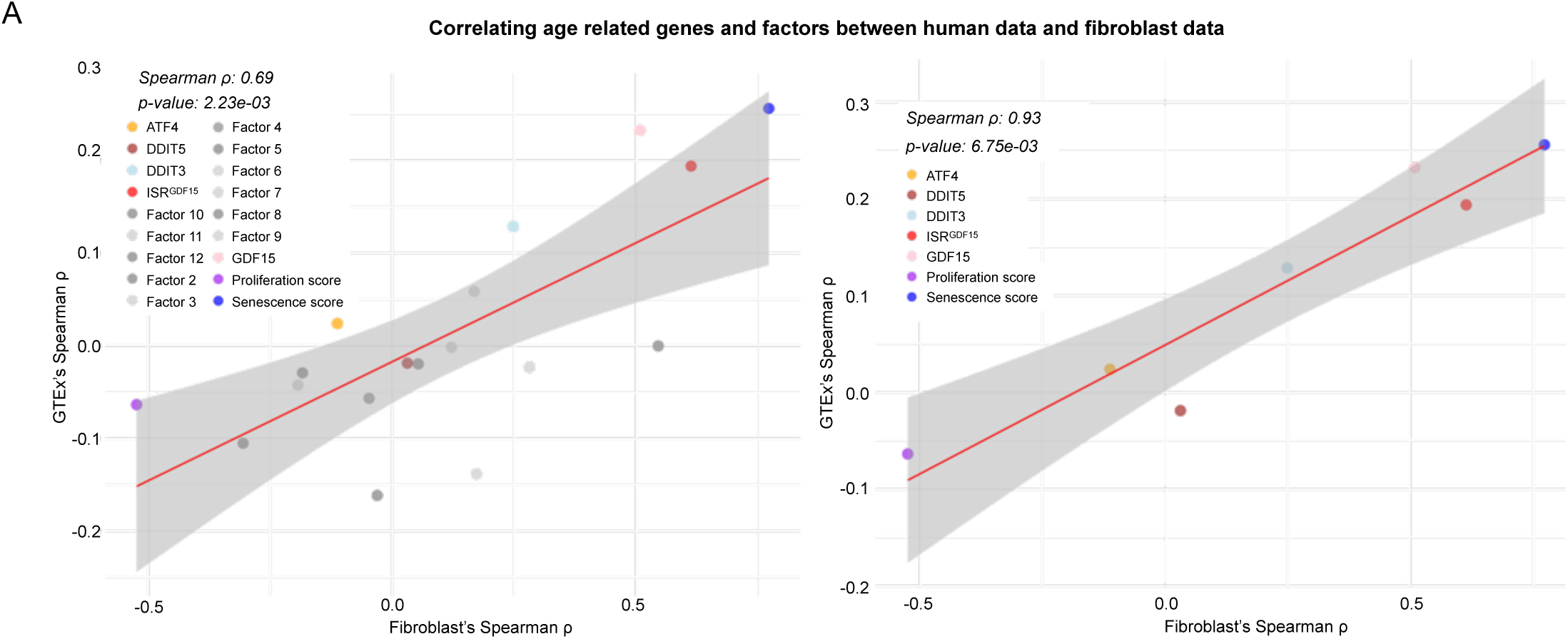
Correlations between human and fibroblast age associations with genes and factors. (**A**) Spearman’s correlations of the average Spearman’s ρs of all tissues comparing donor age to chosen genes, *ISRGDF15* score, or other 11 factor scores (left). The panel on the right shows the same plot without the 11 non-*ISRGDF15* factors.

**Extended Data Figure 8.**
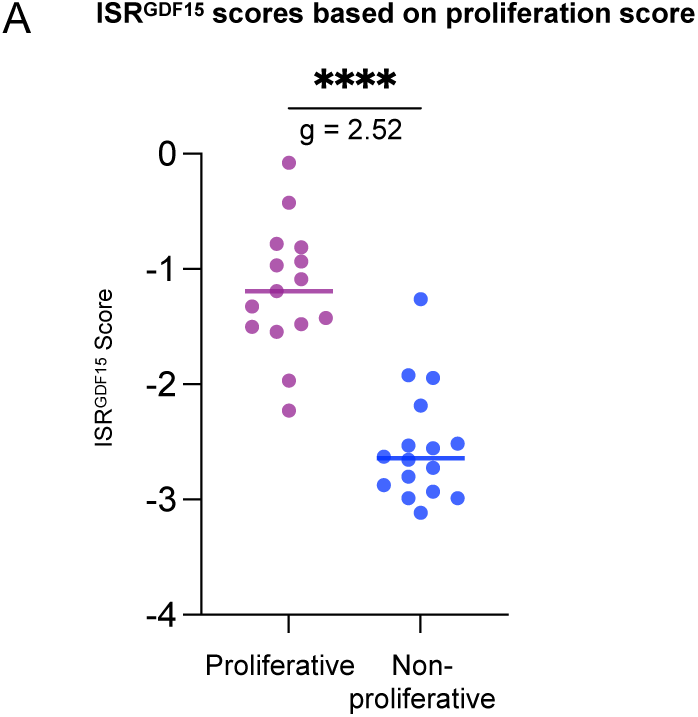
Human tissues’ *ISRGDF15* scores split based on proliferation status. (**A**) *ISRGDF15* score of each tissue split into two groups based on their proliferation score, using the median proliferation score to split the groups. Only the 31 tissues that were significant in Figure 4C are included, p-value determined by Mann-Whitney test.

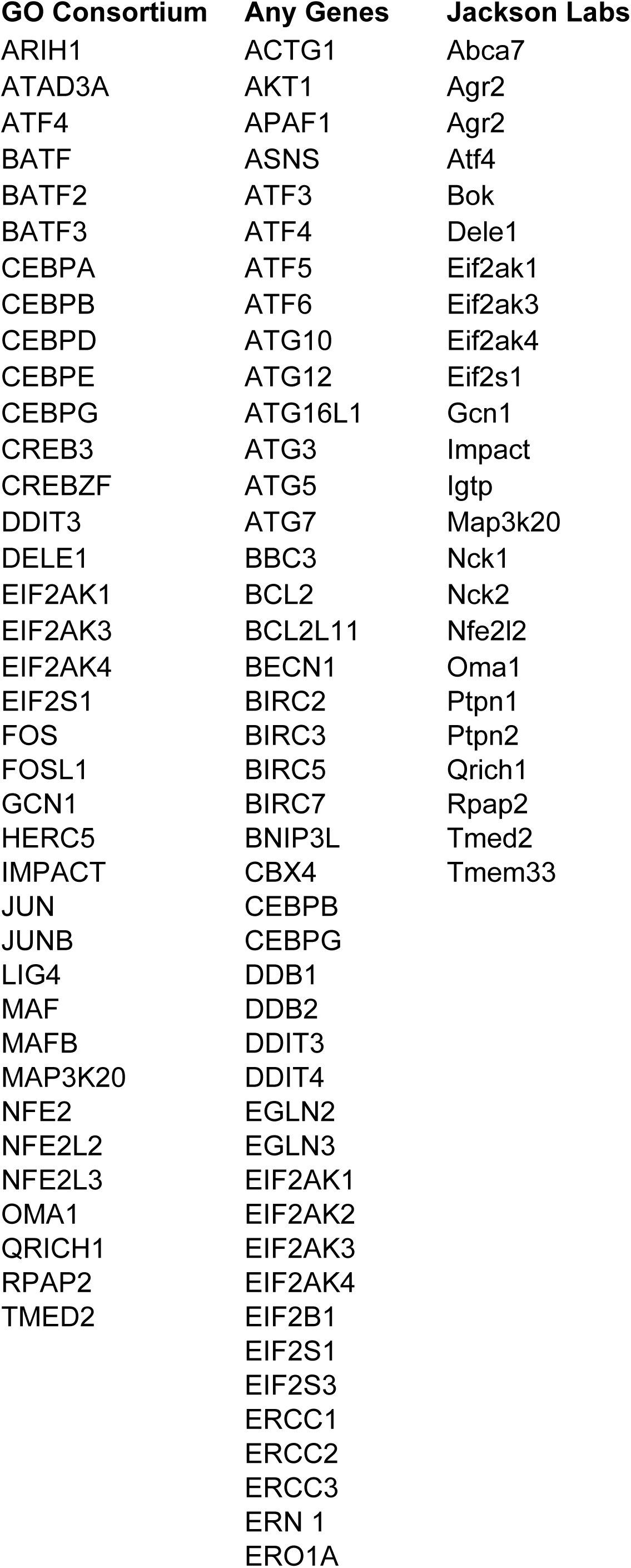
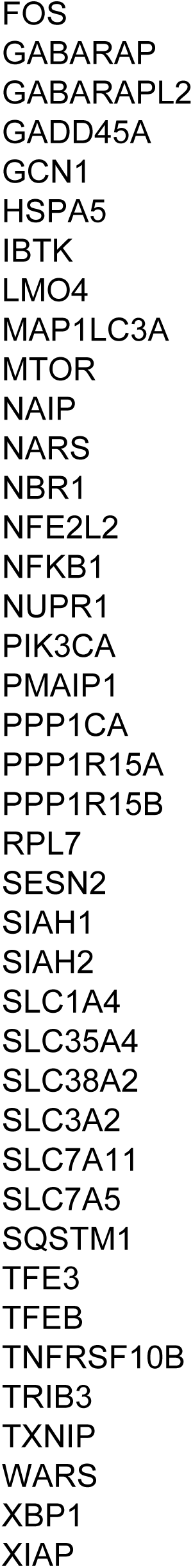
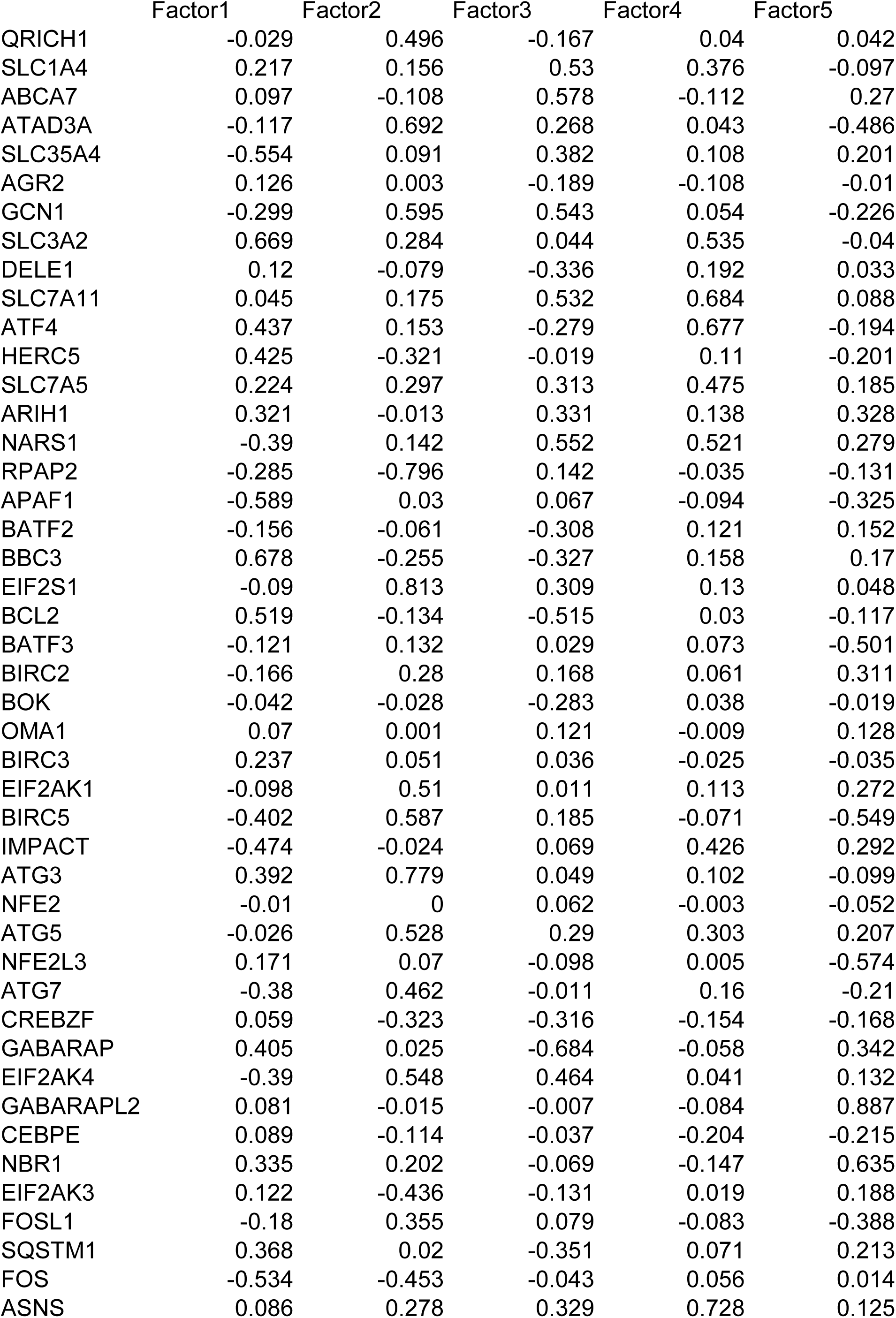
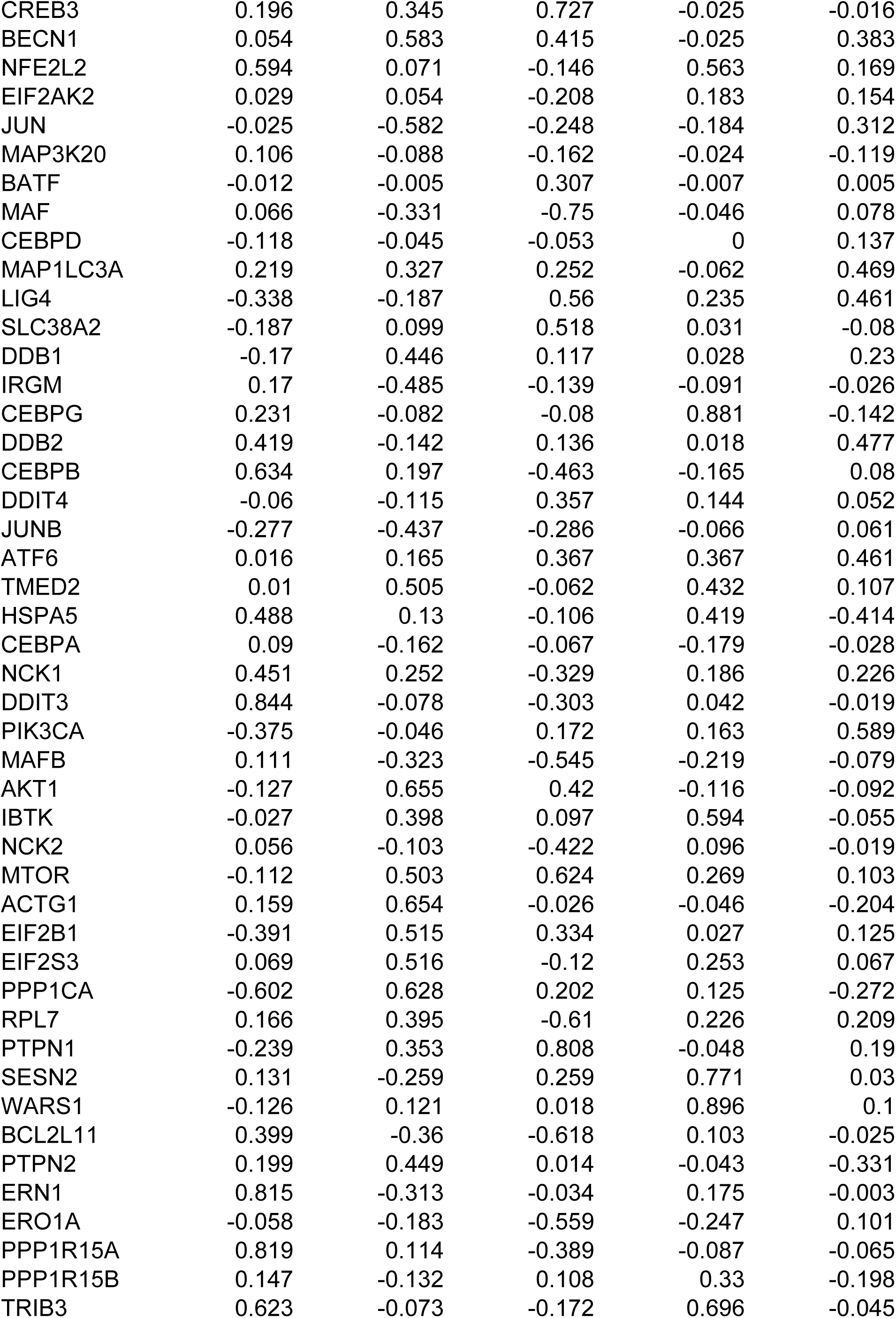
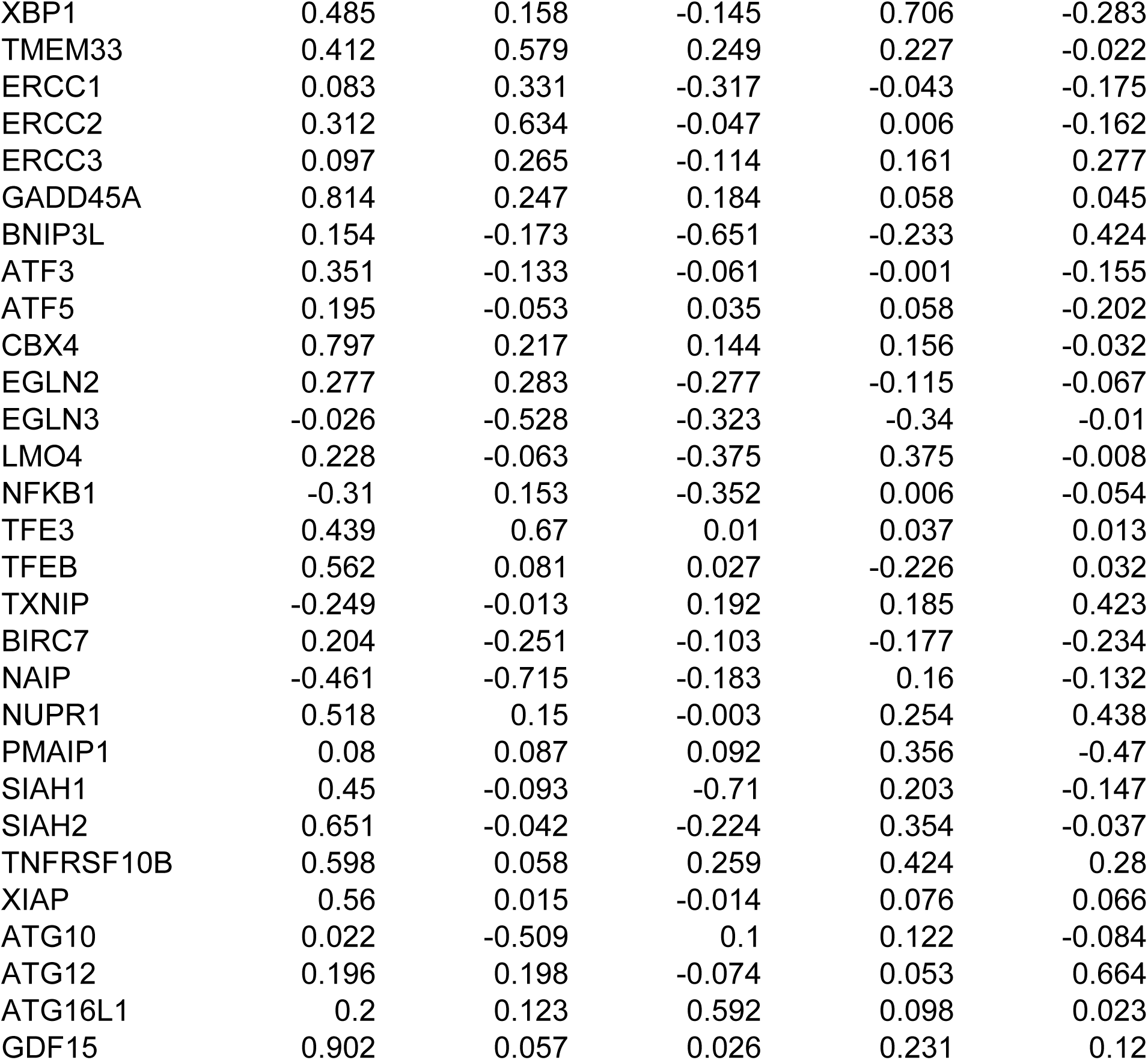
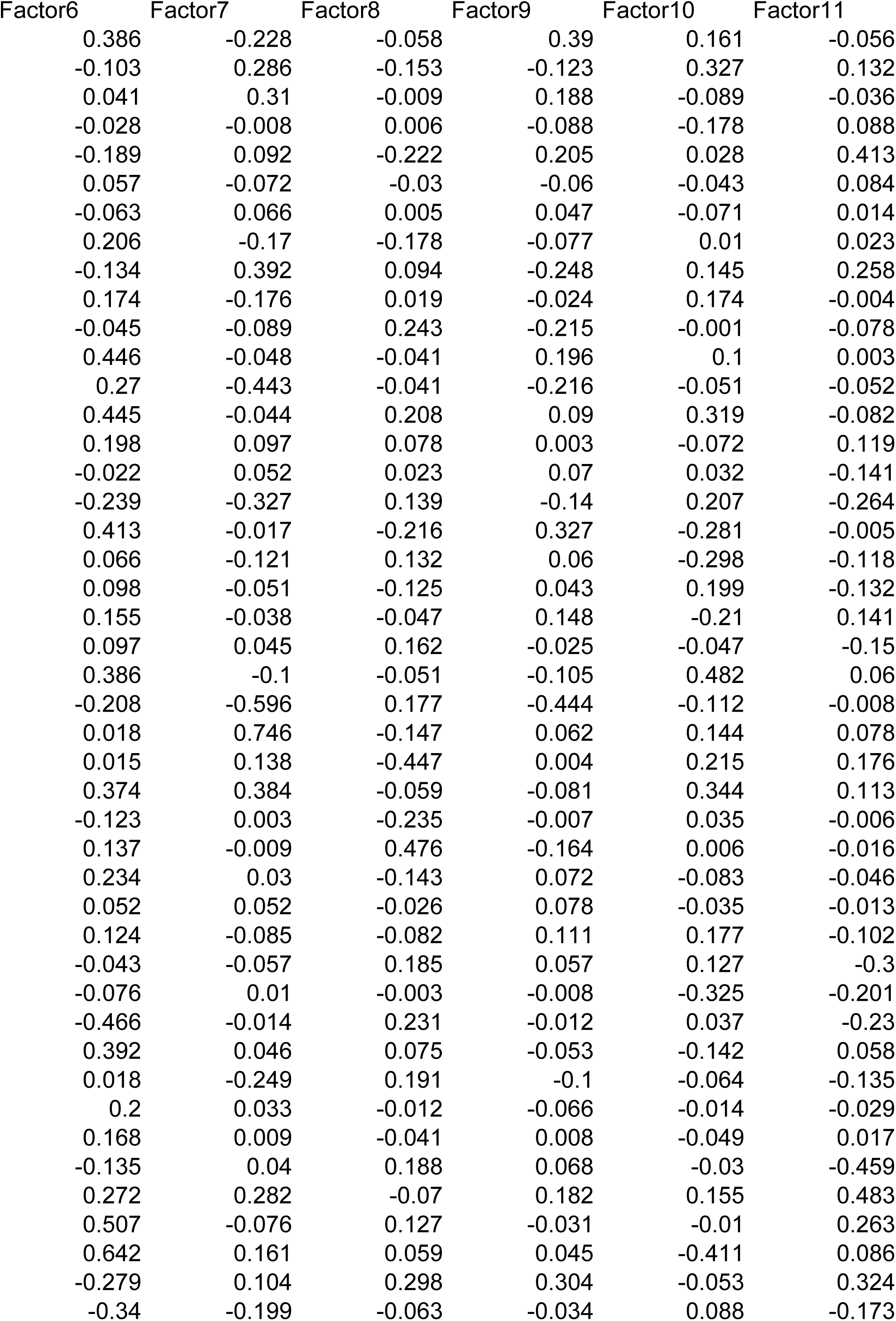
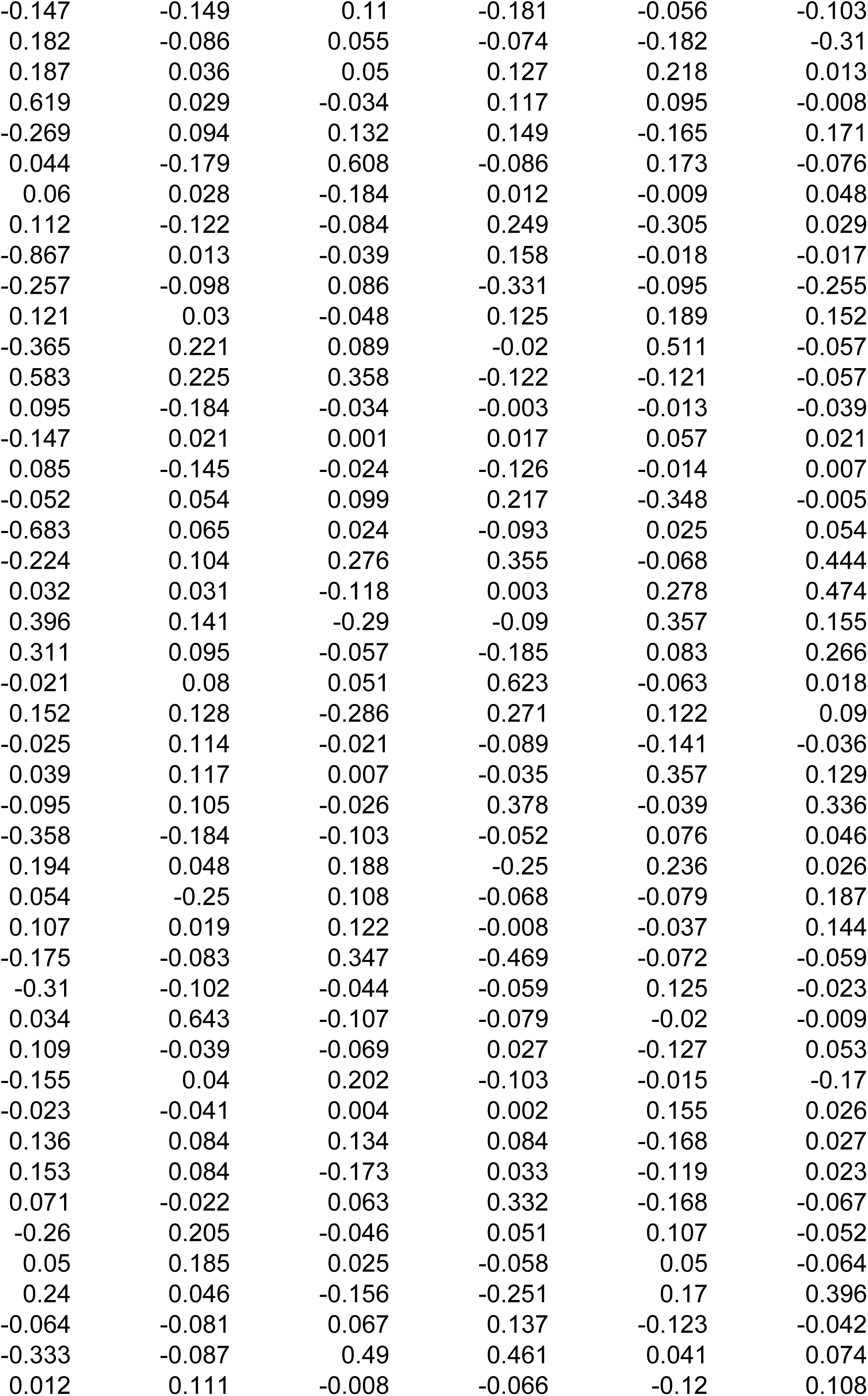
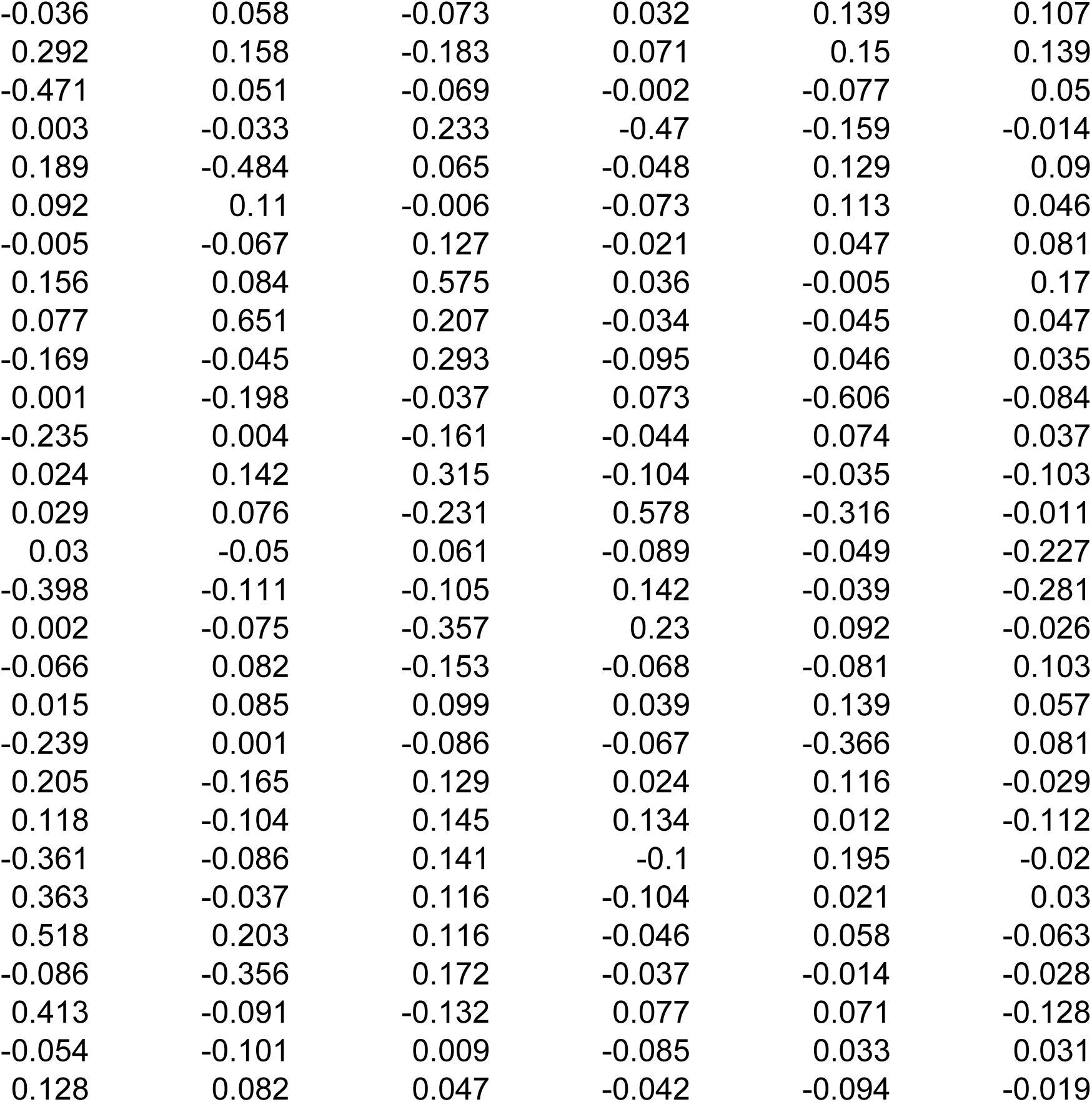
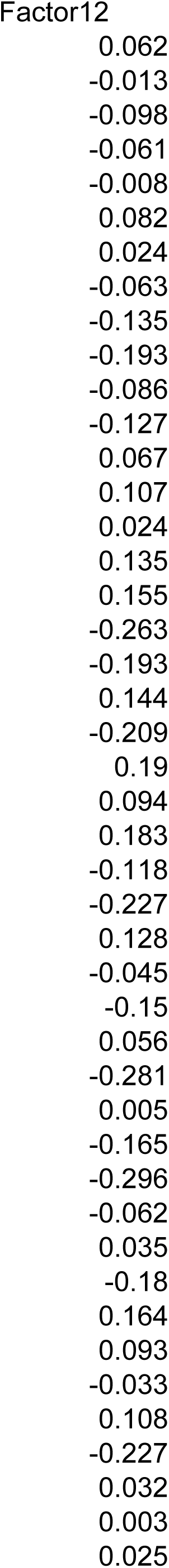
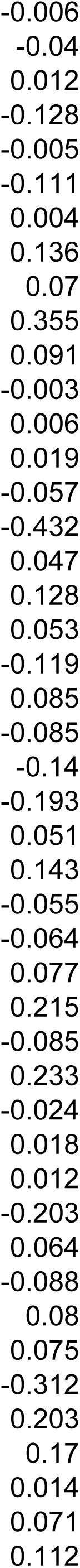
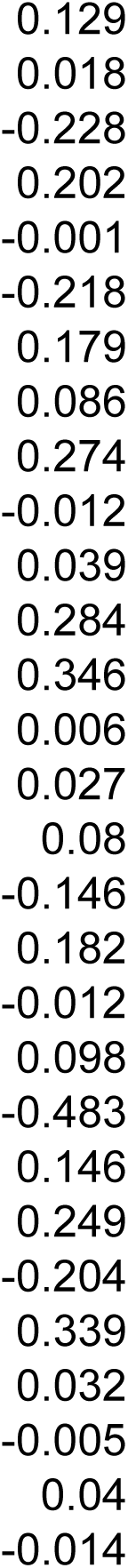
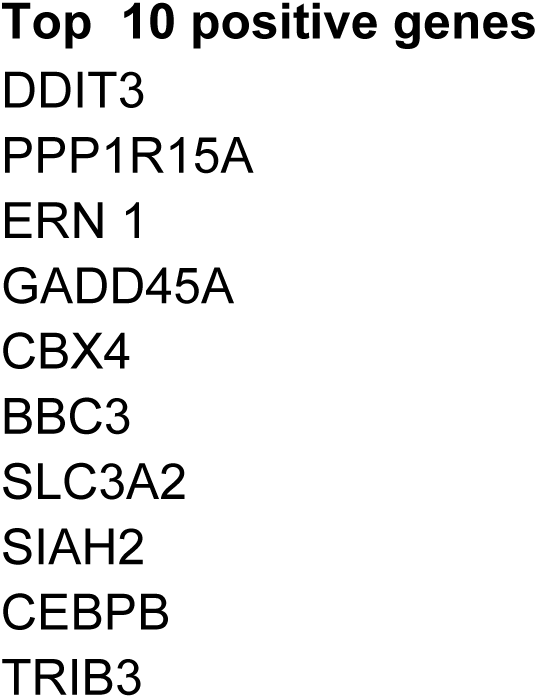
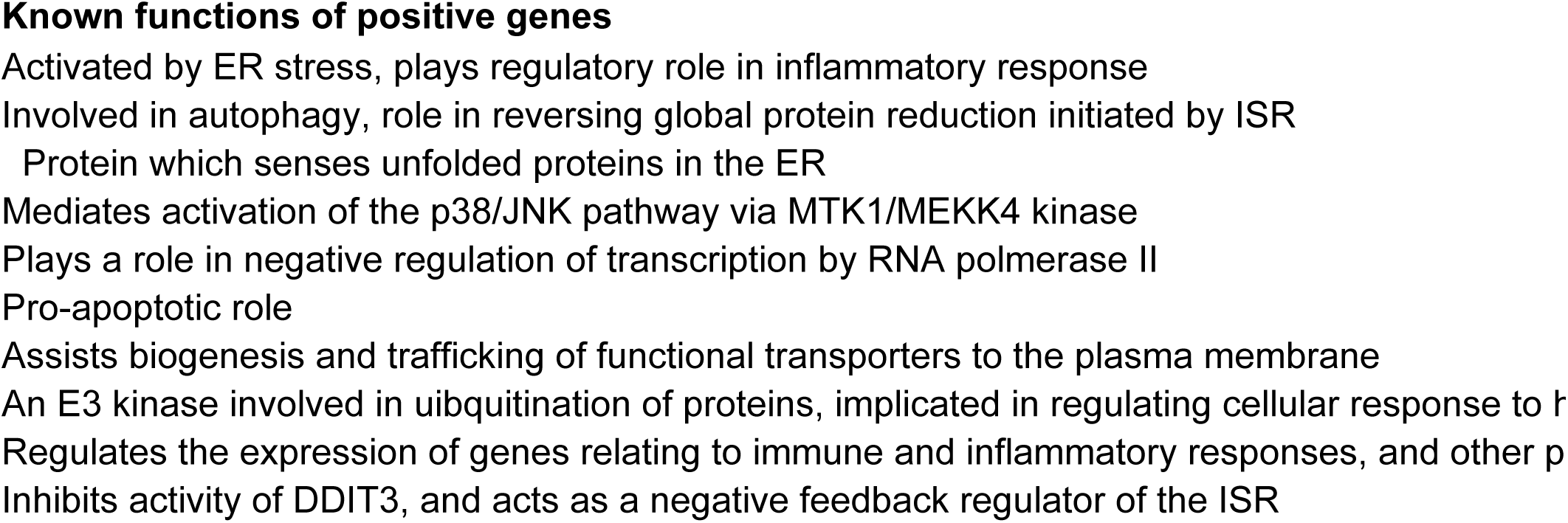
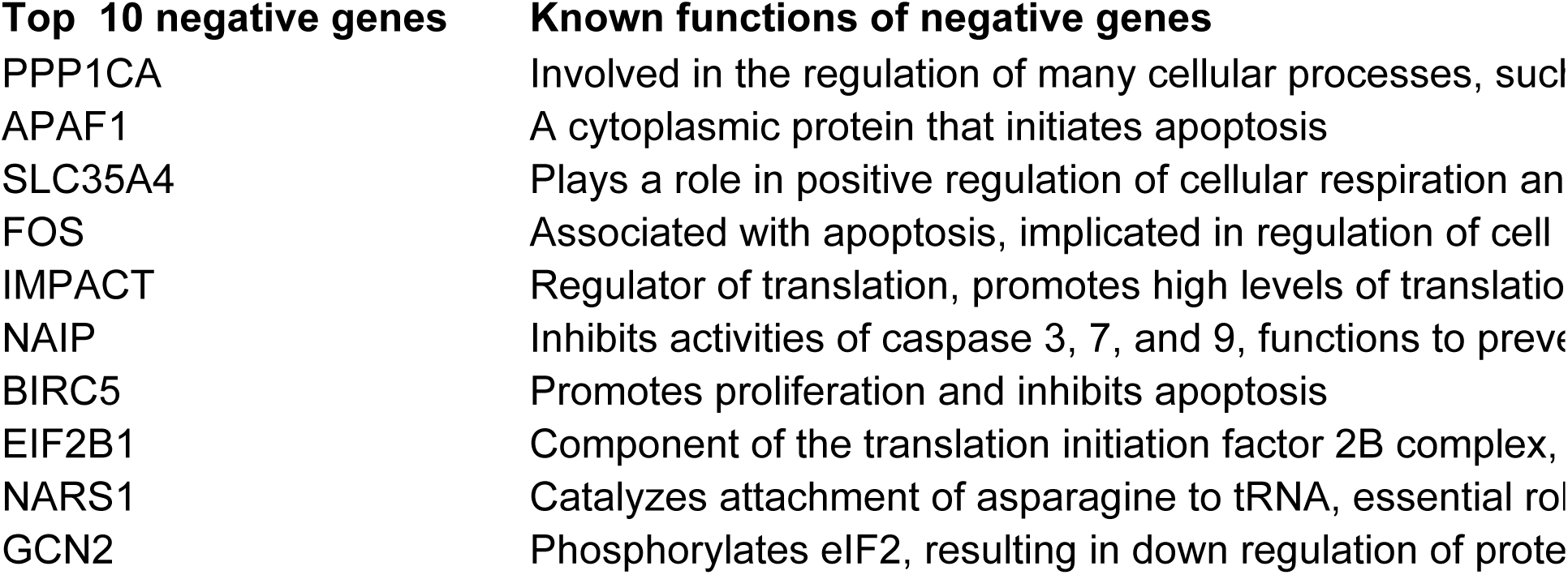
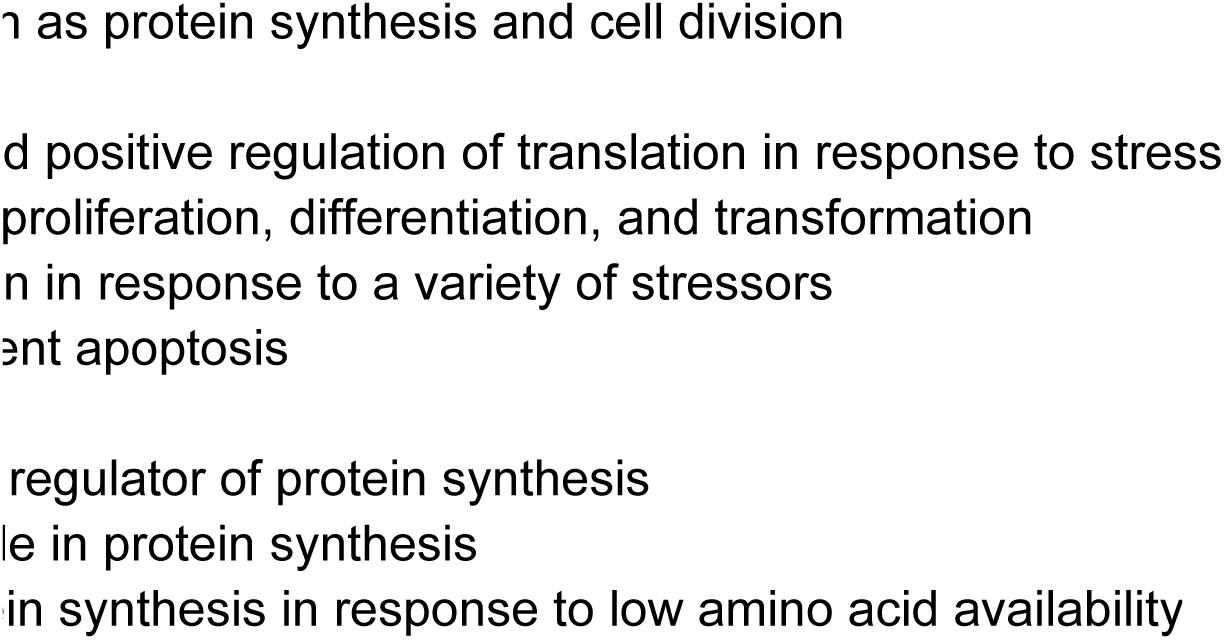
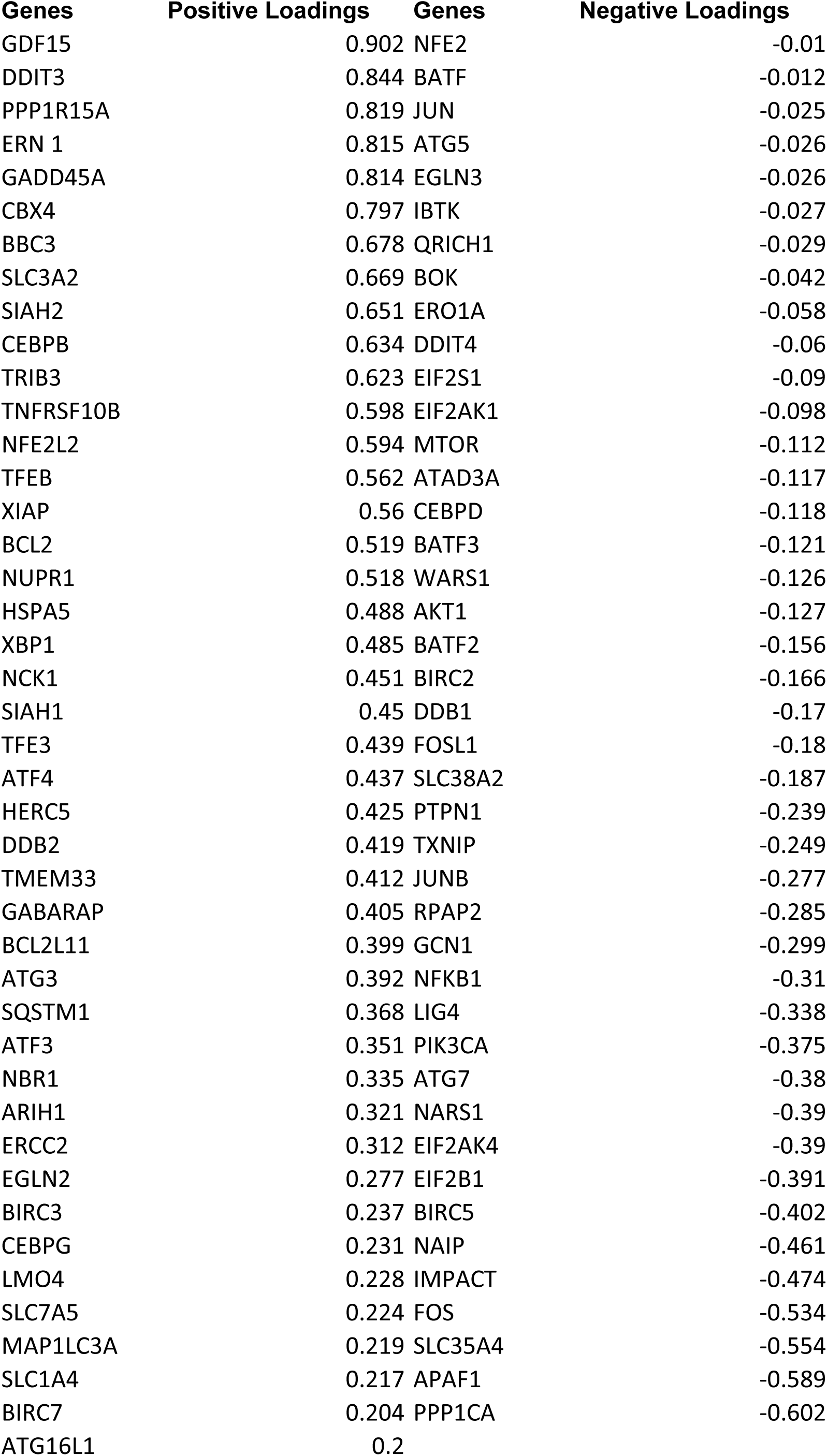
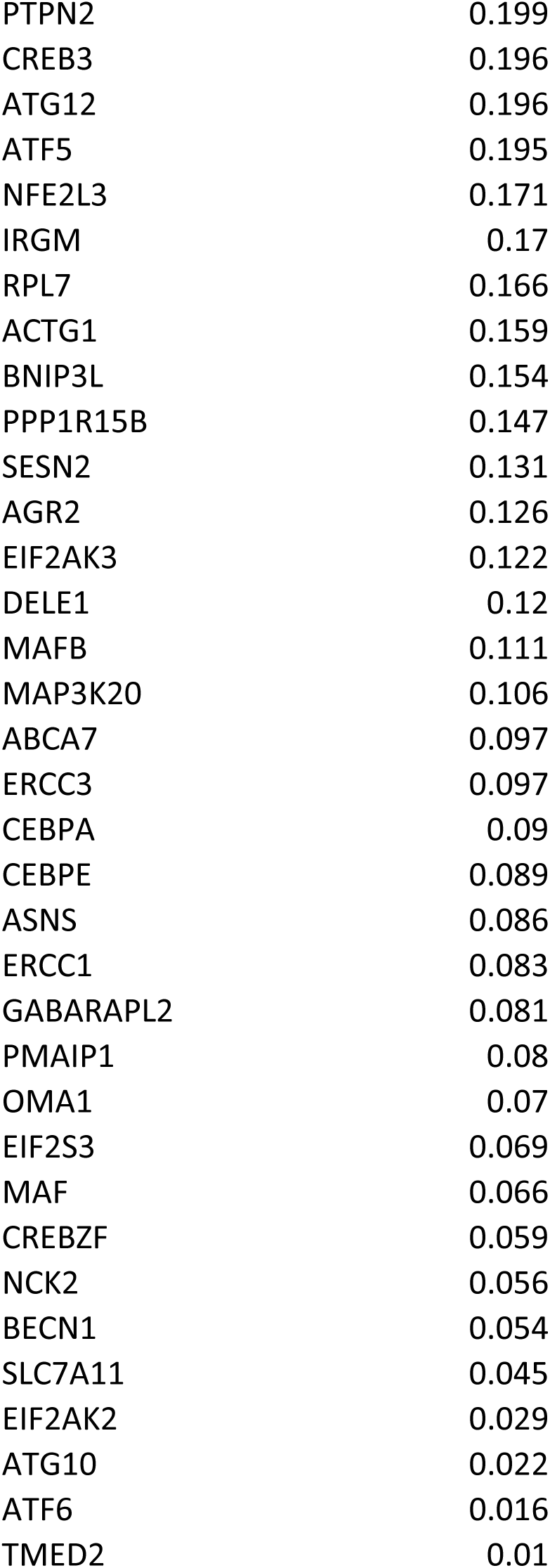

